# Applying stochastic spike train theory for high-accuracy MEG/EEG

**DOI:** 10.1101/532879

**Authors:** Niels Trusbak Haumann, Minna Huotilainen, Peter Vuust, Elvira Brattico

## Abstract

The accuracy of electroencephalography (EEG) and magnetoencephalography (MEG) is challenged by overlapping sources from within the brain. This lack of accuracy is a severe limitation to the possibilities and reliability of modern stimulation protocols in basic research and clinical diagnostics. As a solution, we here introduce a theory of stochastic neuronal spike timing probability densities for describing the large-scale spiking activity in neural networks, and a novel spike density component analysis (SCA) method for isolating specific neural sources. Three studies are conducted based on 564 cases of evoked responses to auditory stimuli from 94 human subjects each measured with 60 EEG electrodes and 306 MEG sensors. In the first study we show that the large-scale spike timing (but not non-encephalographic artifacts) in MEG/EEG waveforms can be modeled with Gaussian probability density functions with high accuracy (median 99.7%-99.9% variance explained), while gamma and sine functions fail describing the MEG and EEG waveforms. In the second study we confirm that SCA can isolate a specific evoked response of interest. Our findings indicate that the mismatch negativity (MMN) response is accurately isolated with SCA, while principal component analysis (PCA) fails supressing interference from overlapping brain activity, e.g. from P3a and alpha waves, and independent component analysis (ICA) distorts the evoked response. Finally, we confirm that SCA accurately reveals inter-individual variation in evoked brain responses, by replicating findings relating individual traits with MMN variations. The findings of this paper suggest that the commonly overlapping neural sources in single-subject or patient data can be more accurately separated by applying the introduced theory of large-scale spike timing and method of SCA in comparison to PCA and ICA.

**Significance statement:** Electroencephalography (EEG) and magnetoencelopraphy (MEG) are among the most applied non-invasive brain recording methods in humans. They are the only methods to measure brain function directly and in time resolutions smaller than seconds. However, in modern research and clinical diagnostics the brain responses of interest cannot be isolated, because of interfering signals of other ongoing brain activity. For the first time, we introduce a theory and method for mathematically describing and isolating overlapping brain signals, which are based on prior intracranial *in vivo* research on brain cells in monkey and human neural networks. Three studies mutually support our theory and suggest that a new level of accuracy in MEG/EEG can achieved by applying the procedures presented in this paper.

## Introduction

### Present limitations in MEG/EEG

Electroencephalography (EEG) and and magnetoencephalography (MEG) methods are among the most applied in human neuroscience (Duncan et al., 2009; Tong and Thakor, 2009). The latest MEG/EEG protocols test advanced cognitive processes and detailed perceptual discrimination abilities for stimuli of increasing complexity (Puce and Hämäläinen, 2017). However, with increasingly complex protocols the neural sources are obtained from fewer measurement samples and show smaller amplitudes compared to other interfering brain activity (Cong et al., 2010). A general problem is that the evoked response of interest becomes difficult to isolate, and the analysis of functional changes in a specific response is often inaccurate and unreliable at the single-subject level (Nikulin et al., 2011; Litvak et al., 2013; Scharf and Nestler, 2018). This leads to low replication rates (Luck and Gaspelin, 2017) and limits the translation of basic MEG/EEG research into clinical applications with the individual patient (Bishop and Hardiman, 2010).

### Isolating the component of interest from a mixture of components

The measured evoked response MEG/EEG waveforms contain a summation of overlapping latent components which must be separated analytically (Luck, 2014). A common solution for isolating a specific component of interest is the use of a functional localizer (Luck and Gaspelin, 2017), or analysis window. The time range and the channel selection for the analysis window is conventionally defined based on the maximum amplitude response in the grand average signal across a group of subjects. A weakness of this method is, however, that other neural sources can remain interfering with the evoked response of interest within the analysis window. Also, a narrow analysis window might result in analytical bias, caused by possible inter-subject variation in the latency and location of the response of interest that may occur outside the analysis window (Luck and Gaspelin, 2017).

### Source location modeling

A more sophisticated popular solution is to separate the overlapping responses by modeling the locations and orientations of the neural sources and their projections through the brain and skull onto the extracranial MEG sensors or EEG electrodes (Wendel et al., 2009). However, when more sources are simultaneously present and the signal-to-noise and interference ratio (SNIR) (including the interference from spatially and temporally overlapping neural activity originating from different brain regions) is low, source location errors of up to centimeters and distorted source waveforms are commonly observed (Schwartz et al., 1999; Vanrumste et al., 2001; Whittingstall et al., 2003; Sharon et al., 2007; Kiebel et al., 2008; Zumer et al., 2008). Recently, it has been considered that a major contribution to the source location modeling errors may originate from the simultaneous estimation of the source amplitudes, locations, orientations and projections within a single model (Wendel et al., 2009). Instead, it has been considered to first separate the mixed sources with blind source separation, prior to modeling the locations and orientations of the sources (Vigario et al., 2000; Zhukov et al., 2000; Richards, 2004; Tsai et al., 2006; Reynolds and Richards, 2009).

### Blind source separation

With blind source separation it is assumed that each component has a consistent spatial distribution, or topography. The component topography is represented by a linear weighting vector that defines the magnitude and polarity of the projection of the component waveform onto each MEG/EEG channel, which is often estimated with principal component analysis (PCA) or independent component analysis (ICA) (Choi et al., 2005). However, a general weakness of the blind source separation methods is that they cannot separate sources with similar spatial topographies, and they do not distinguish between sources based on their polarity (Groppe et al., 2008). These are nevertheless two crucial characteristics for identifying sources originating from the brain (Picton et al., 1974). Therefore, we here suggest applying a novel spike density component analysis (SCA) method, which in addition to the spatial topography also models the polarity and temporal shape of the neural sources of extracranial MEG/EEG measurements reflecting the large-scale spiking activity constituted by individual spike timing behavior of the neurons in brain networks.

### Large-scale stochastic neuronal spike trains

The electrical potentials measured with EEG and the magnetic fields measured with MEG originate from large-scale spiking activity of neurons and the resulting postsynaptic potentials in neural networks (Hämäläinen et al., 1993; Deco et al., 2008; Buzsaki et al., 2012). Each spike of the single neuron involves an action potential in the axon of typical 1 ms duration and a synaptic current flow with a duration in the range of 10 ms, which typically results in a local voltage change of 25 mV and magnetic field of 20 fAm across 0.1-0.2 mm (Hämäläinen et al., 1993). The spiking activity observed in intracranial recordings of the electrophysiological responses to auditory, visual or tactile stimuli of single cortical or subcortical neurons is commonly analyzed with a peristimulus time histogram (PSTH) (Rodieck, 1962; Gerstein and Mandelbrot, 1964; Dorrscheidt, 1981; deCharms and Merzenich, 1996; Brown et al., 2004; Filali et al., 2004; Shimazaki and Shinomoto, 2007; Mukamel et al., 2010). The PSTH shows the number of spikes counted in time bins, i.e., the momentary firing rate in spikes per second, in a time window locked to the presentation of a stimulus. The spikes in each time bin are counted across a series of trials of repeated stimulation. Whereas the spike timing of the single neuron after each single stimulation appears to be random, the accumulated spike timing across a series of trials reveals systematic distributions of the spikes across time, which can be described with stochastic spike density functions (Rodieck, 1962; Gerstein and Mandelbrot, 1964; Barbieri et al., 2001; Brown et al., 2004; Stein et al., 2005; Maimon and Assad, 2009; Teramae and Fukai, 2014).

The spike densities, observed as variance in the spike timing of the single neuron, have been considered to originate from a large-scale principle of ‘stochastic resonance’ in neural networks, which depends on the organization of the synaptic pathways (Stein et al., 2005; Teramae and Fukai, 2014). While the spike timing variability in single neurons is commonly described with stochastic functions (Gerstein and Mandelbrot, 1964; Barbieri et al., 2001; Shin, 2002; Stein et al., 2005; Maimon and Assad, 2009; Teramae and Fukai, 2014; Aljadeff et al., 2016), it has not yet been investigated how the stochastic variance in spike timing might be reflected in EEG and MEG. PSTHs for peripheral neurons show regular clock-like spike patterns with low variability in the spike timing, such as in neurons in the brain stem, while in pyramidal cortical neurons, in particular in association areas, there is higher variability in spike timing, as observed with intracranial single neuron recordings (Maimon and Assad, 2009). Interestingly, non-invasive scalp EEG recordings of evoked responses from the human brain stem reveal similar narrow time distributions of each response component (I, II, III, IV, V, VI), while the cortical evoked responses (N1, P2, N2) observed from cortical regions exhibit similar broader temporal distributions (Picton et al., 1974). Based on these considerations, we suggest that, in addition to single-neuron spike timing behaviour, also large-scale neuronal activity in MEG/EEG waveforms might be systematically described with stochastic spike density functions (Figure 1).

**Figure 1.**
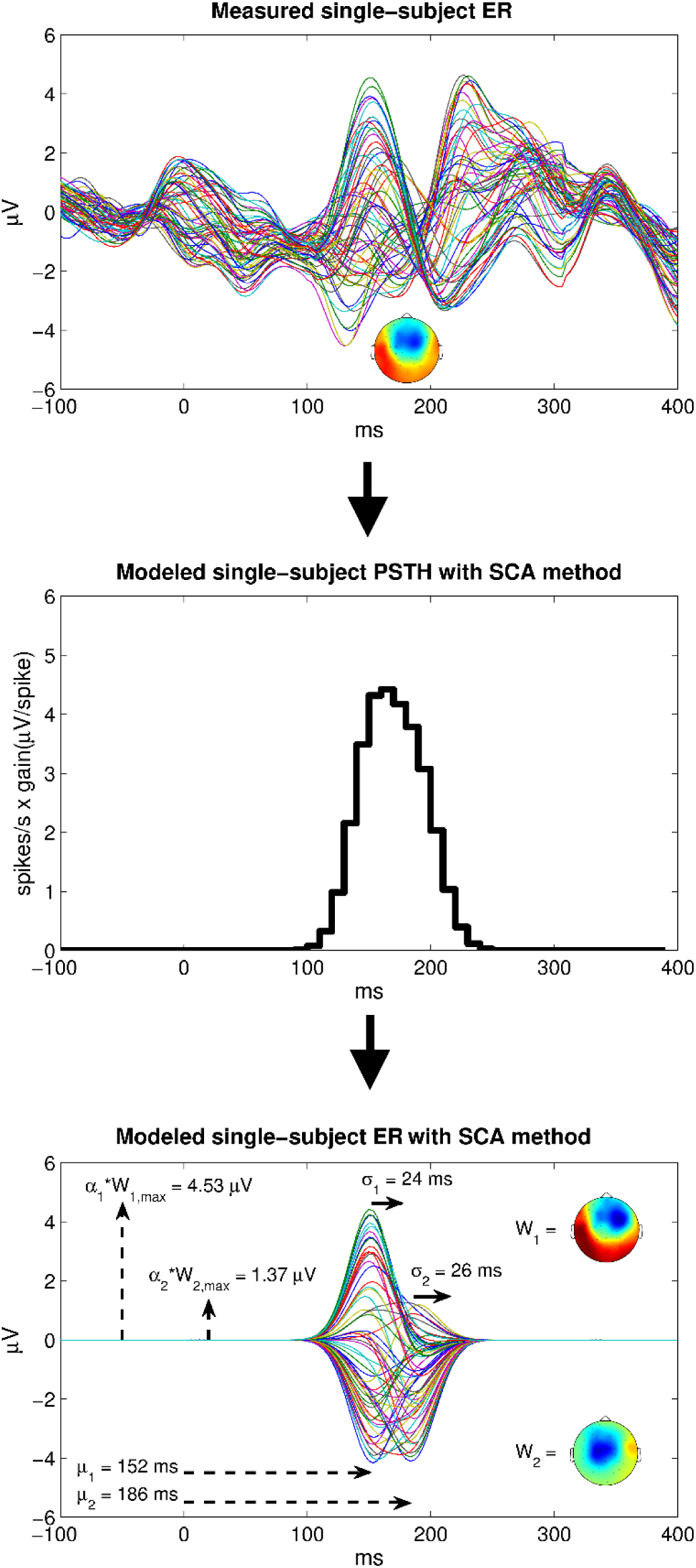
Modeling single-subject EEG waveforms with stochastic spike density functions. In the top is shown an example of a measured single-subject evoked response (ER). The middle depicts a reinterpretation of the same data modeled as large-scale PSTH with the SCA method. In the bottom is shown the parameters for the data modeled with Gaussian density functions.

The main generators of the spiking activity observed in MEG/EEG waveforms are cortical pyramidal neurons from layers IV-V (Hämäläinen et al., 1993; Friston, 2005). At the micrometer scale of the single neuron, the spike timing of the cortical pyramidal cell has often been described as a Poisson process (Gerstein and Mandelbrot, 1964; Barbieri et al., 2001; Kass et al., 2003; Stein et al., 2005; Maimon and Assad, 2009; Waldert et al., 2013; Teramae and Fukai, 2014; Aljadeff et al., 2016). At the centimeter scale of electrocorticography (ECoG), EEG and MEG a large number of spikes are involved in generating the spike density (Hämäläinen et al., 1993), and Poisson processes with large numbers of events can be approximated by Gaussian probability density functions (Tseng, 1949). Therefore, we suggest that MEG/EEG waveforms can be modeled by Gaussian functions (see Methods section, Formula 1).

### Research questions and hypotheses

In Study 1 we investigated whether single-subject average evoked responses (ERs) measured with EEG and MEG can be modeled by stochastic functions. The modeling performance of Gaussian functions was measured in percent explained variance and compared to the modeling performance of gamma and sine functions. Also, we investigated whether Gaussian functions can specifically model the signals originating from the brain or other signals such as eye blink artifacts. This was tested by comparing the modeling performance and the residual signal peak amplitudes for Gaussian functions either with preprocessing or without preprocessing, in the last case including artifactual signals not originating from the brain.

In our second study we tested whether the spike density component analysis (SCA) based on the Gaussian functions can be applied to accurately isolate a specific evoked brain response of interest, such as the mismatch negativity (MMN) response. Moreover, we investigated whether SCA isolates the MMN more accurately compared to the commonly applied principal component analysis (PCA) and independent component analysis (ICA). The accuracy of the MMN extraction was measured based on the number of MMN-related components (fewer components indicates higher accuracy), the correlations between the extracted single-subject MMN response and the group-level MMN with respect to its spatial topography and temporal morphology (higher correlations indicate higher accuracy), and the root-mean-squared error (RMSE) of the remaining interfering signals in the baseline time points surrounding the MMN response (the lower RMSE the higher accuracy).

Furthermore, in our third study we tested whether previous findings of individual differences in MMN amplitude related to depressive traits analyzed using conventional functional localizers (Bonetti et al., 2017) could be replicated with the novel SCA method.

## Results of Study 1

The SCA decomposition method was tested on a database with 564 cases of average auditory evoked responses of healthy adult subjects measured simultaneously with 60 EEG electrodes, 102 axial MEG magnetometer sensors and 204 planar MEG gradiometer sensors.

An example of typical result of an SCA decomposition with Gaussian functions is shown in Figure 2. Significant differences are observed in the percent explained variance of SCA with Gaussian functions compared to gamma or sine functions, χ^2^(2)=867.14, *p*<10^−188^ (Figure 3). Post-hoc comparisons show that the Gaussian function outperforms the gamma function (*p*<10^−93^) and sine function (*p*<10^−93^) (Figure 3 and Table 1). Also, the sine function performs slightly better than the gamma function (*p*<10^−11^) (Figure 3 and Table 1). In general, the SCA with the Gaussian function show gradual increases in the explained variance by the components, while the SCA modeled with gamma and sine functions fails explaining more variance after the first component estimate for the peak amplitude in the MEG/EEG waveform (Figure 4).

**Table 1.** Percent explained variance by Gaussian, gamma, and sine functions. Post hoc comparisons of percent explained variance in EEG, MEG magnetometer (MAG) and MEG gradiometer (GRAD) after decomposing the single-subject evoked responses into Gaussian, gamma and sine components.

|  |  | Median performance in<br>% explained variance | Post-hoc comparisons, *p* |  |
| --- | --- | --- | --- | --- |
| SCA function | Modality |  |  |  |
| Gaussian | EEG | 99.9 | EEG | MEG mag. |
|  | MEG mag. | 99.7 | <10<sup>-37</sup> * |  |
|  | MEG grad. | 99.7 | <10<sup>-38</sup> * | .217 |
| Gamma | EEG | 25.3 | - | - |
|  | MEG mag. | 20.9 | - | - |
|  | MEG grad. | 20.1 | - | - |
| Sine | EEG | 26.6 | - | - |
|  | MEG mag. | 21.6 | - | - |
|  | MEG grad. | 21.5 | - | - |

**Figure 2.**
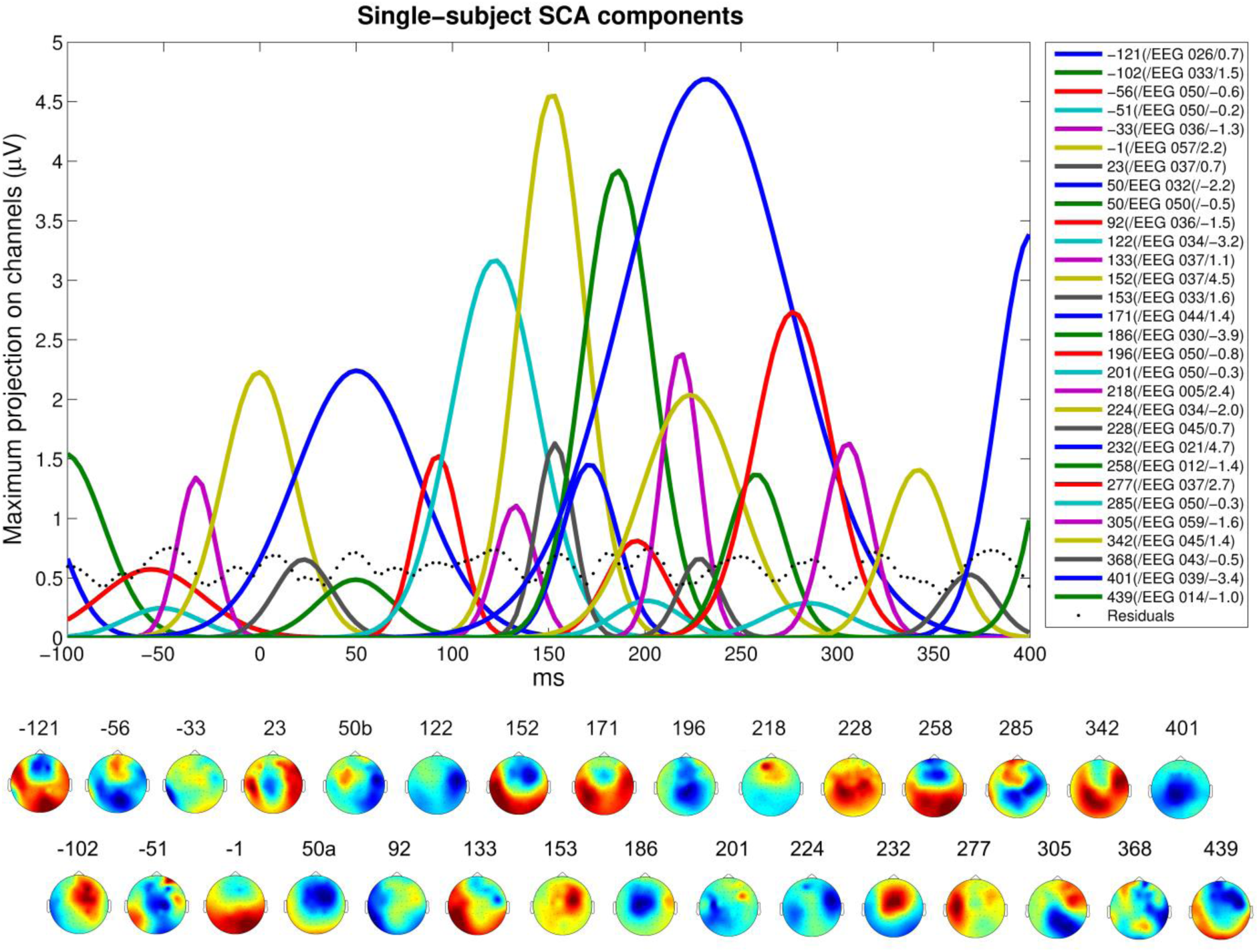
An example of a single-subject waveform decomposed into SCA components. SCA components for a single-subject and stimulus condition (slide deviant) with the peristimulus time in ms on the horizontal axis and the EEG amplitude of the SCA components in the peak channel in μV on the vertical axis (irrespective of differences in peak channels across components). Below is shown the topographies of the components (color scales are set according to the maximum amplitude for each component). Numbers shown next to the component labels (right) and topographies (bottom) designates expected latency in ms. Component labels (right) are defined by expected latency in ms, name of peak channel, and amplitude (here in μV).

**Figure 3.**
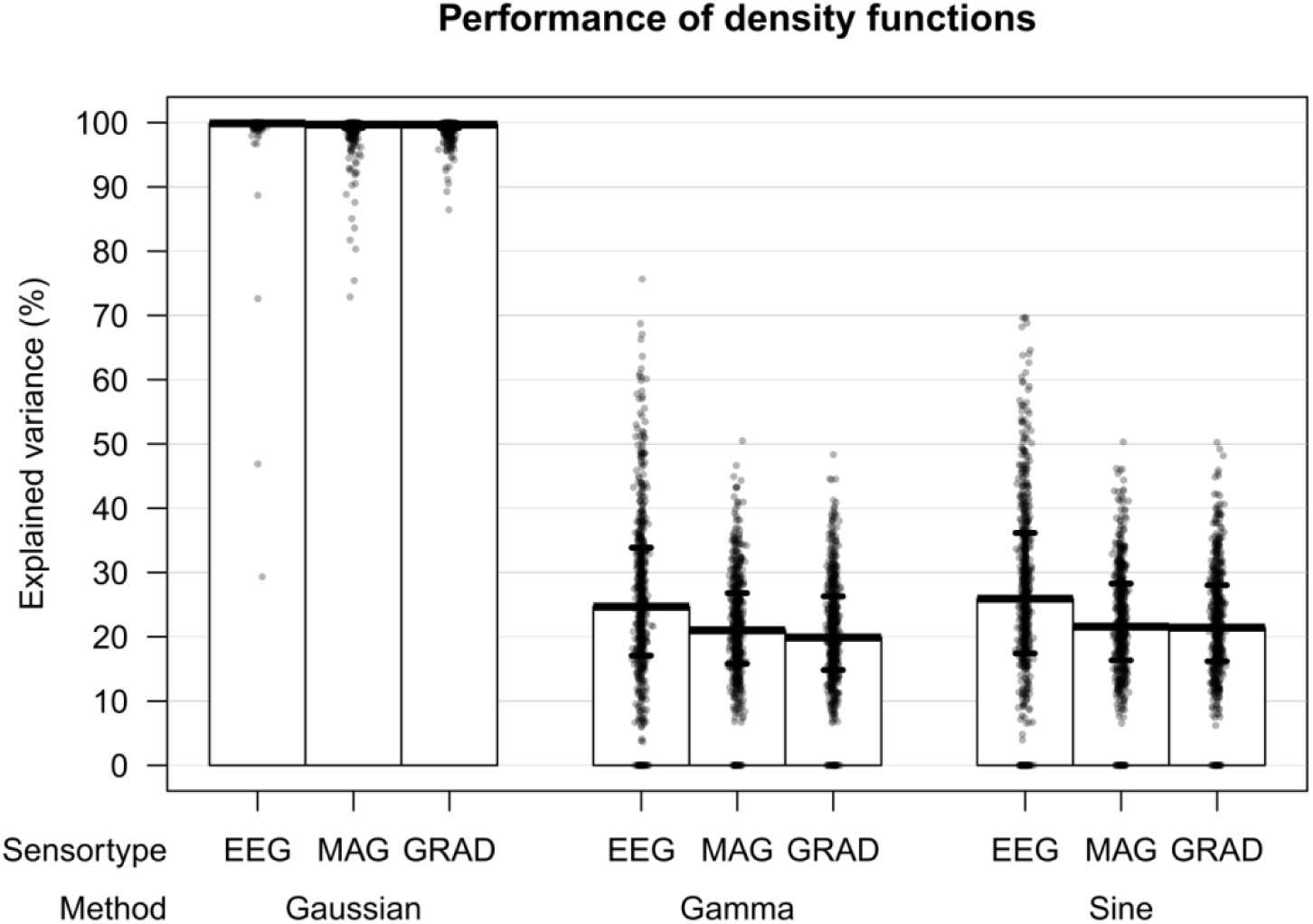
Percent explained variance in MEG/EEG waveforms by Gaussian, gamma, and sine functions. Showing the percent explained variance in EEG, MEG magnetometer (MAG) and MEG gradiometer (GRAD) after decomposing the single-subject evoked responses into Gaussian, gamma and sine components.

**Figure 4.**
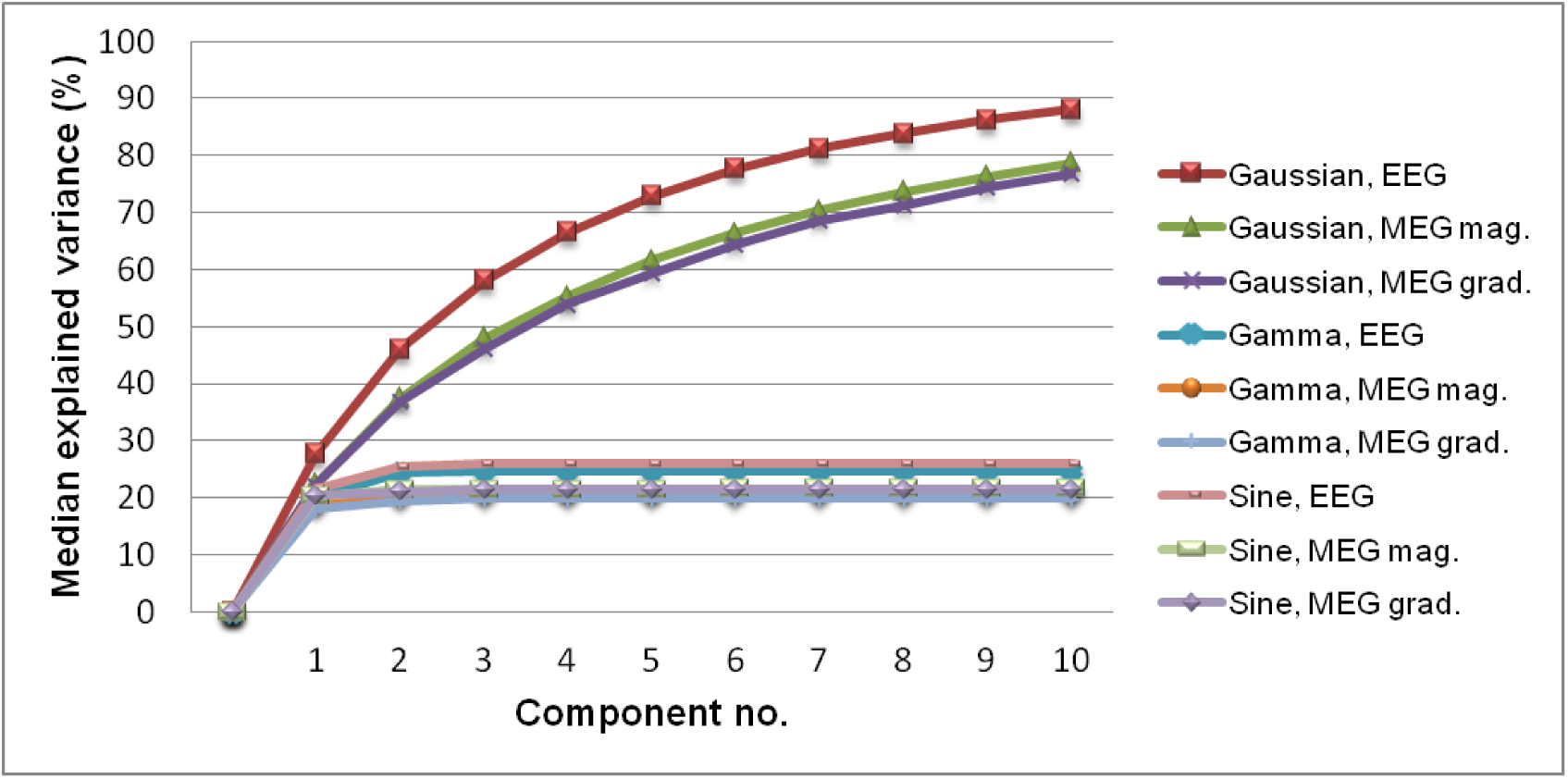
Cumulative percent variance explained by the first ten Gaussian, gamma, and sine components. Showing the accumulated percent variance explained in EEG, MEG magnetometer (MAG) and MEG gradiometer (GRAD) after decomposing the single-subject evoked responses into the first ten Gaussian, gamma and sine components of highest amplitudes in descending order of amplitude.

Across measurement modalities, the Gaussian function shows a slightly higher modeling performance on the EEG waveforms compared to the MEG waveforms, while there is no significant difference in the Gaussian modeling performance on the MEG magnetometer and gradiometer waveforms (Figure 3 and Table 1, and Figure 4). Interestingly, the shape parameter, *k*, of the gamma function shows the highest median value in the EEG (*k*=48.8), and gradually lower values for the MEG magnetometers (*k*=38.7), and MEG gradiometers (*k*=30.1), χ^2^(2)=26.49, *p*<10^−5^, indicating increasing skewness 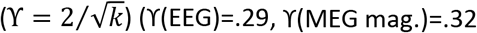, and ϒ(MEG grad.)=.37) dependent on the measurement modality.

There is significant decrease in the modeling performance with Gaussian functions on the average MEG/EEG waveforms that have not been preprocessed compared to those that have been preprocessed, *p*<10^−90^ (Figure 5). Also, the peak amplitudes in the residual waveforms is higher for the Gaussian SCA models without preprocessing compared to with preprocessing in the EEG, *p*<10^−64^, MEG magnetometers, *p*<10^−24^, and MEG gradiometers, *p*<10^−49^ (Figure 6). The grand average waveforms obtained with all the compared methods are shown in Figure 7.

**Figure 5.**
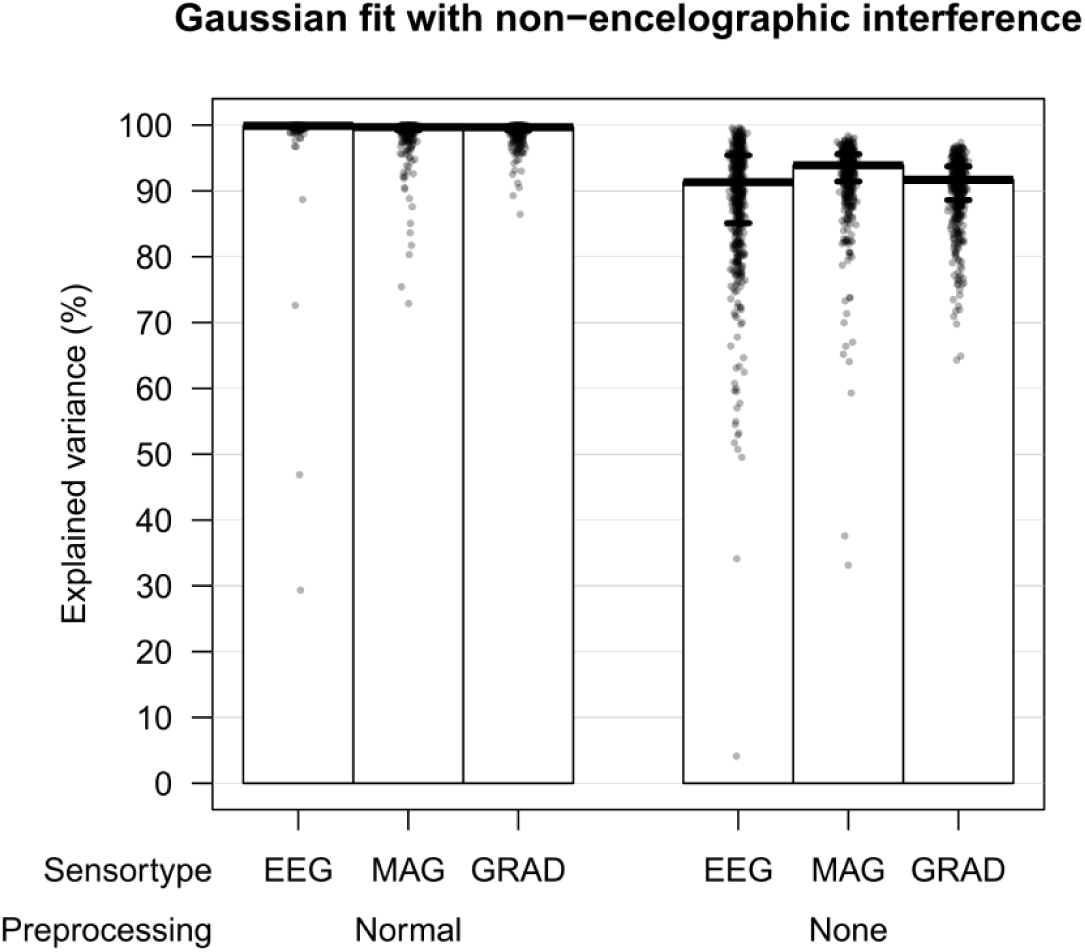
Percent explained variance with (normal) and without (none) preprocessing to remove artifacts. Showing the percent explained variance in EEG, MEG magnetometer (MAG) and MEG gradiometer (GRAD) after decomposing the single-subject evoked responses with normal preprocessing or no (none) preprocessing into Gaussian components.

**Figure 6.**
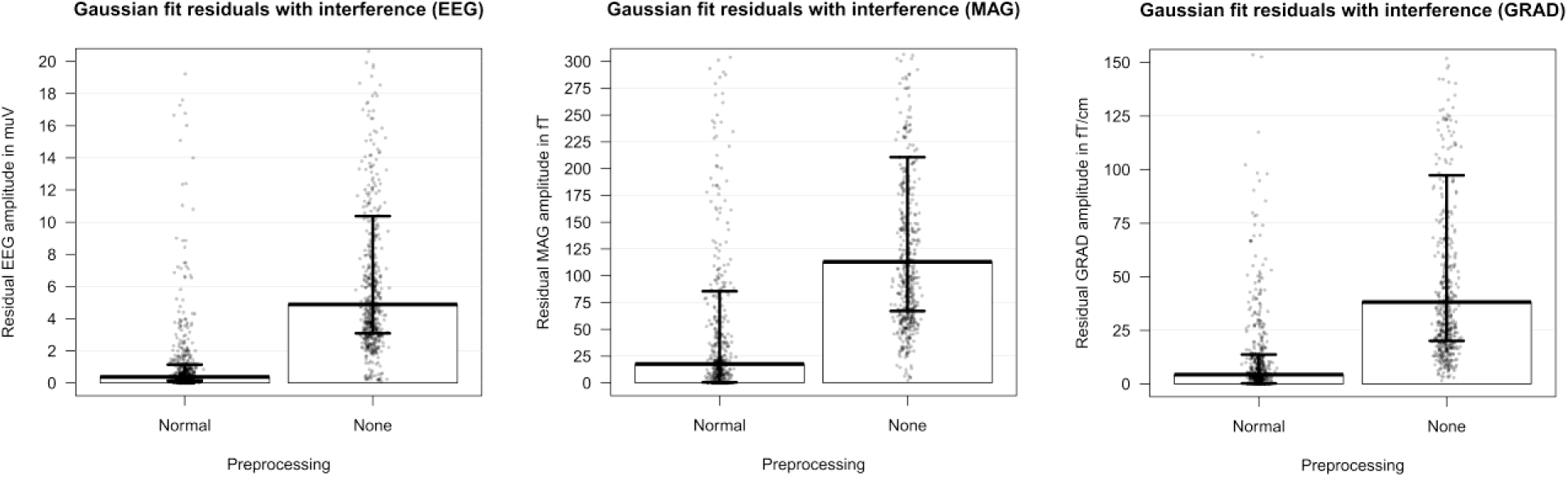
Residual peak amplitudes with (normal) and without (none) preprocessing to remove artifacts. Showing peak amplitudes of the remaining variance in the EEG, MEG magnetometer (MAG) and MEG gradiometer (GRAD) waveforms which could not be decomposed into Gaussian components.

**Figure 7.**
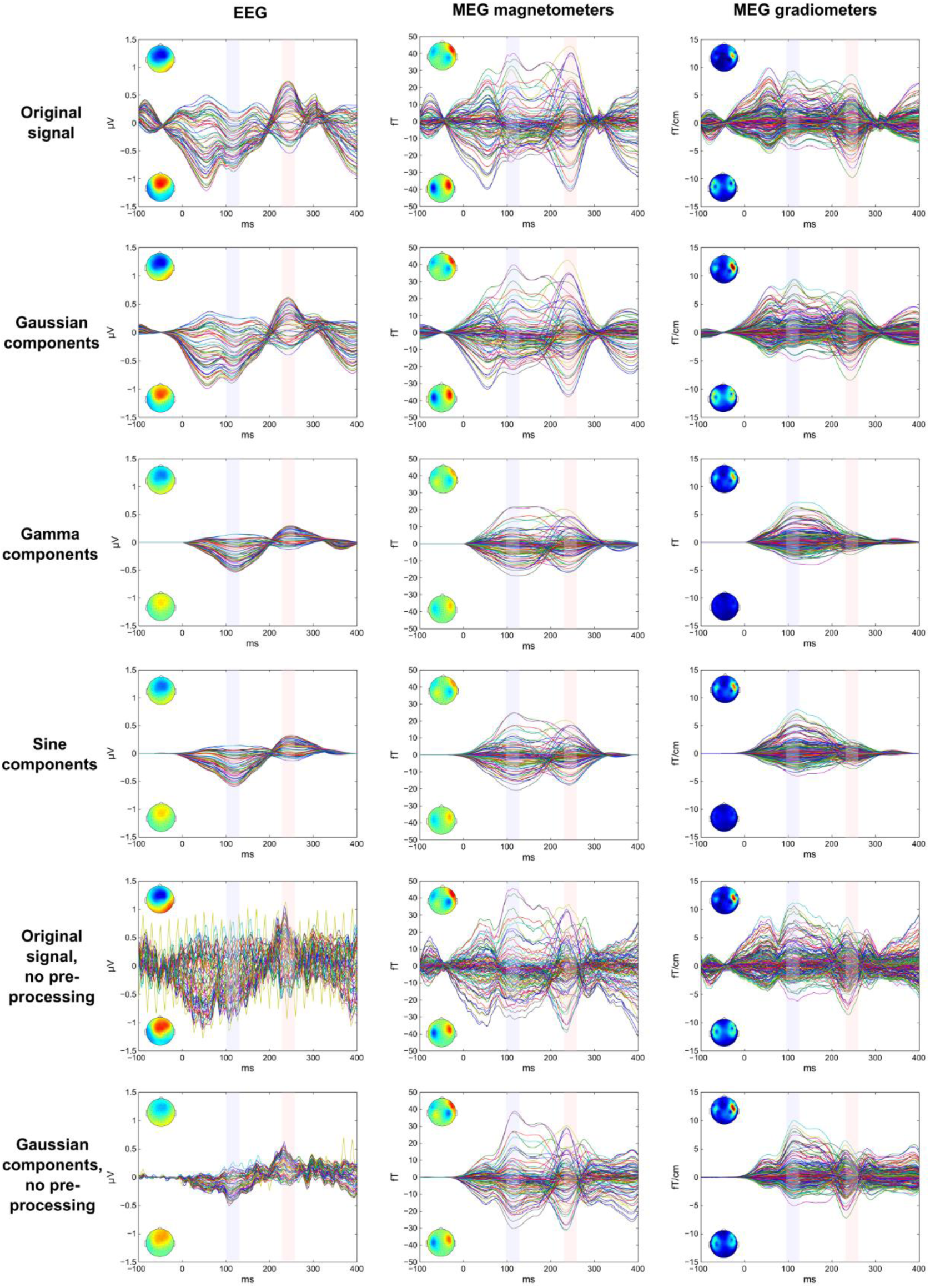
Components extracted with Gaussian, gamma and sine functions across all cases. Showing the grand-average butterfly waveform plots for the extracted first five SCA components across all 564 cases. Next to each waveform is shown the topography by using a 30 ms time window around the peak latencies of the negative component (MMN) marked with a light blue rectangle (peak latencies: EEG=116 ms, MEG magnetometers=113 ms, MEG gradiometers=110 ms) and the positive component (P3a) marked with a light red rectangle (peak latencies: EEG=243 ms, MEG magnetometers=245 ms, MEG gradiometers=246 ms).

## Interim discussion 1

The findings in the first study support our hypothesis that large-scale spike density components in MEG/EEG waveforms originating from neural networks can accurately be described by Gaussian functions, with median 99.7%-99.9% of the variance explained by Gaussian functions. While the first component, of highest amplitude, to some extend could be approximated by gamma and sine functions, our findings suggest that the Gaussian function better models the complete set of spike density components in the MEG/EEG waveforms. It seems unlikely that the high performance of the Gaussian function is related to the bandpass filter response, because the bandpass filter has a constant response shape, while the modeled Gaussian components vary in width, and the performance of the Gaussian function is still relatively high without the preprocessing with bandpass filtering excluded. As with any MEG/EEG analysis in general, with the SCA method it is necessary to consider a compromise between the suppression of artifactual signals while retaining as much of the component of interest across frequency bands. In addition, it is unlikely that the Gaussian distribution is caused by timing error in the signal averaging across trials, because the component of interest in the single-trial evoked responses is typically identical in shape in the average single-subject waveform when the stimulus condition remain constant (Gaspar et al., 2011). Moreover, while it might be considered that the current sources of the spike density moves in space as it propagates on the cortex, findings from research on ‘micro-states’ suggests that the component topography does not change significantly across time except in the transitions between components (Lehmann, 1989; Pascualmarqui et al., 1995; Koenig et al., 2014), which supports the validity of applying constant channel (*c*) weights, *W*_*n,c*_, for each component (*n*). The overlap between components when their relative amplitudes changes will result in topographies that through visual inspection appear to be moving across time, even though the topography of each separate component might be relatively constant.

Interestingly, the findings also showed that the skewness defined by the gamma shape parameter was lowest for EEG, higher for MEG magnetometers, and highest for MEG gradiometers. This might reflect that the signal gain in the measurement angle of the EEG electrodes causes less overlap between the MMN and P3a responses of opposite polarities, while this overlap has a stronger influence on the signal gain of the MEG magnetometer and gradiometer sensors. Another explanation could be that the higher skewness and the slightly lower percent explained variance in MEG compared to EEG is related to general differences in the spatial specificity affecting how much of the complete spiking distribution is included in the measurements. The measured part of the large-scale spike timing distribution in a neural network might be most complete and thus most symmetrically distributed in the largest-scale EEG measurements, less complete and thus more skewed in the more spatially specific MEG magnetometers, more incomplete in the additionally spatially specific MEG gradiometers, and most incomplete in the highly spatially specific intracranial measurements that show the highest skewness in the measured spike timing distribution (Maimon and Assad, 2009).

While the findings in Study 1 suggests that MEG/EEG waveforms can accurately be decomposed into spike density components with the SCA method, in the following Study 2, we investigated whether the SCA method can be applied to isolate a specific evoked response of interest from spatially and temporally overlapping neural sources with higher accuracy compared to PCA and ICA.

## Results of Study 2

The same dataset as in Study 1 was applied to compare evoked response decompositions with SCA, ICA and PCA. An automatic template matching method was used to extract the MMN-related components.

We observed a significant difference in the number of components extracted from the SCA, ICA, and PCA decompositions representing the MMN, χ^2^(2)=10.49, *p*=.005 (Figure 8). Post-hoc comparisons showed that there was in general fewer components representing the MMN in the SCA decompositions compared to the ICA (*p*<10^−4^) and PCA (*p*<10^−4^) decompositions, while ICA and PCA decompositions tended to contain similarly large numbers of components representing the MMN (*p*=.167). The SCA decompositions contained different numbers of components representing the MMN dependent on the measurement modality, χ^2^(2)=342.63, *p*<10^−74^ (Figure 8). The SCA decompositions of the MEG gradiometer waveforms contained slightly more (maximum = 6) components representing the MMN compared to those for the MEG magnetometer waveforms (maximum = 2) (*p*<10^−39^) and EEG waveforms (maximum = 2) (*p*<10^−59^).

**Figure 8.**
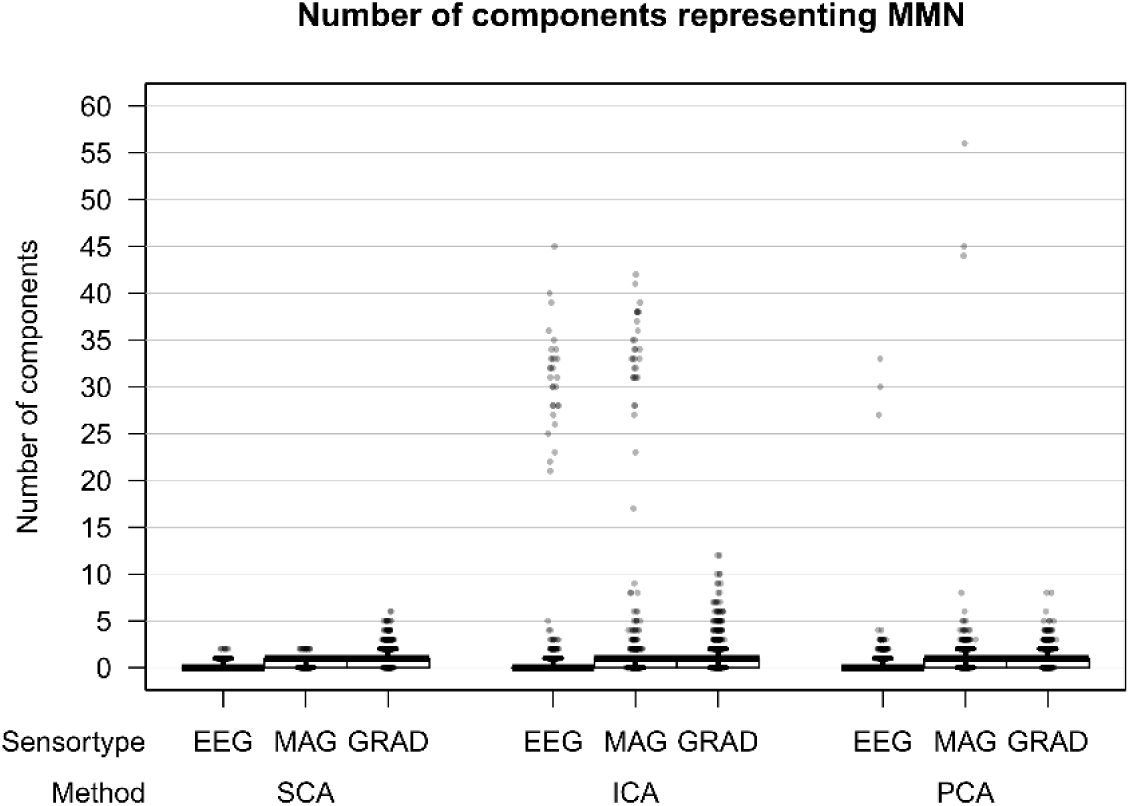
Number of SCA, ICA and PCA components representing the MMN. Showing number of SCA, ICA and PCA components representing MMN based on EEG, MEG magnetometer (MAG) and MEG gradiometer (GRAD) waveforms.

The similarity of the single-subject and group MMN topography differed significantly between SCA, ICA, PCA and the original, χ^2^(3)=162.80, *p*<10^−34^ (Figure 9). All component analysis methods resulted in more accurate representations of the MMN topography compared to the original (Figure 9 and Table 2). While the ICA components represented the MMN topography slightly more accurately compared to the SCA and PCA components, similar performances in accuracy of topography were observed for the SCA and PCA components (Figure 9 and Table 2). Moreover, the MMN topography was more accurately represented by the SCA components for EEG than for MEG magnetometers (*p*<10^−21^) and MEG gradiometers (*p*<10^−26^), and for MEG magnetometers more accurately than for MEG gradiometers (*p*=.015) (Figure 9).

**Table 2.** Similarity of topography and waveform between single-subject and group MMN. Post hoc comparisons on topography and waveform similarities for EEG, MEG magnetometer (MAG) and MEG gradiometer (GRAD) waveforms applying SCA, ICA, PCA or the original waveforms.

| Similarity between topography of single-subject and group MMN |  |  |  |  |
| --- | --- | --- | --- | --- |
| Method | Median performance in $r^2$ | Post-hoc comparisons, $p$ | | |
| SCA | .49 | SCA | ICA | PCA |
| ICA | .51 | $<10^{-5} *$ | | |
| PCA | .47 | .296 | $<.001 *$ | |
| Original | .35 | $<10^{-20} *$ | $<.10^{-36} *$ | $<10^{-30} *$ |
| Similarity between waveform of single-subject and group MMN |  |  |  |  |
| Method | Median performance in $r^2$ | Post-hoc comparisons, $p$ | | |
| SCA | .47 | SCA | ICA | PCA |
| ICA | .31 | $<10^{-64} *$ | | |
| PCA | .42 | .005 * | $<10^{-57} *$ | |
| Original | .35 | $<10^{-47} *$ | $<10^{-33} *$ | $<10^{-39} *$ |

**Figure 9.**
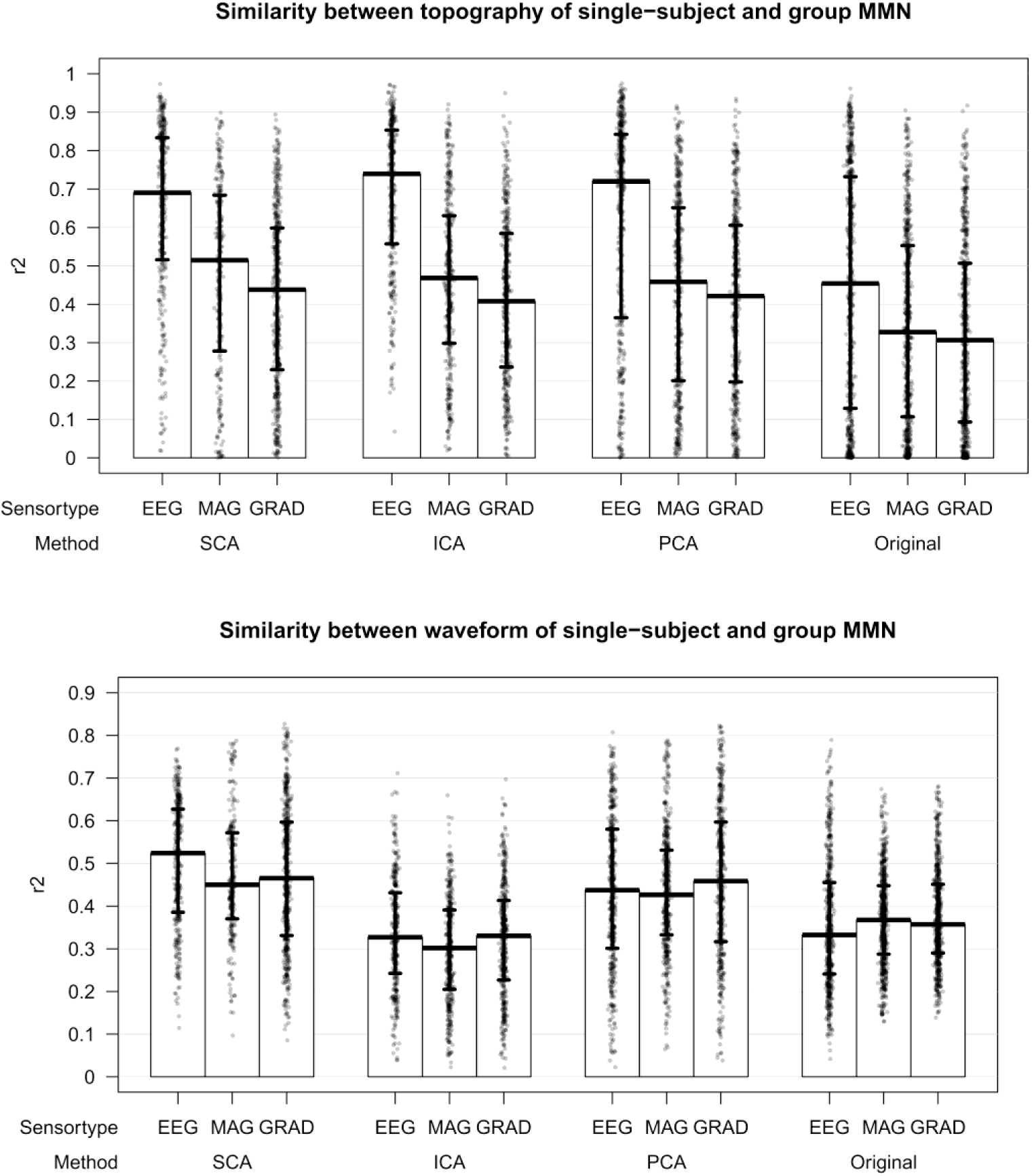
Similarity of topography and waveform between single-subject and group MMN. Showing topography and waveform similarities for EEG, MEG magnetometer (MAG) and MEG gradiometer (GRAD) waveforms applying SCA, ICA, PCA or the original waveforms.

The similarity between the extracted single-subject MMN components and the group-level MMN differed significantly dependent on the applied method, χ^2^(3)=543.23, *p*<10^−116^ (Figure 9). The extracted single-subject SCA components showed the highest similarity with the group-level MMN waveforms (Figure 9 and Table 2), while also the PCA components showed higher resemblance with the group-level MMN waveforms compared to the original single-subject waveforms. However, the representation of the MMN waveform with the ICA components was worse than the original single-subject waveforms (Figure 9 and Table 2). There was a minor decrease in the performance on representing the MMN waveform with SCA components for the MEG magnetometers compared to the EEG (*p*<10^−5^) and MEG gradiometers (*p*<10^−9^), while a similar high performance was observed for the EEG and the MEG gradiometers (*p*=.728).

Also, with respect to the removal of the interfering signals overlapping with the MMN components, the SCA method outperformed the other methods for the EEG, χ^2^(3)=1193.77, *p*<10^−257^, MEG magnetometers, χ^2^(3)=1149.56, *p*<10^−248^, and gradiometers, χ^2^(3)=1026.73, *p*<10^−221^ (Figure 10, Table 3 and Figure 11). The best removal of interfering signals was achieved with the SCA, followed by the ICA, and PCA. The resulting grand averages of the MMN topographies and waveforms achieved with each component analysis method is shown in Figure 12.

**Table 3.** Remaining interfering signals overlapping with component isolated with SCA, ICA, and PCA. Post hoc comparisons for the root-mean-squared error (RMSE) from a perfect MEG/EEG waveform baseline of values 0 in the time range surrounding the component of interest, *t*_*comp*_, for EEG, MEG magnetometer (MAG) and MEG gradiometer (GRAD) waveforms when applying SCA, ICA, PCA or the original waveforms.

| Remaining interfering signals (EEG) |  |  |  |  |
| --- | --- | --- | --- | --- |
| Method | Median $\mu V$ | Post-hoc comparisons, $p$ | | |
| SCA | 0.002 | SCA | ICA | PCA |
| ICA | 0.072 | $<10^{-25} *$ | | |
| PCA | 0.407 | $<10^{-65} *$ | $<10^{-41} *$ | |
| Original | 1.164 | $<10^{-92} *$ | $<10^{-92} *$ | $<10^{-86} *$ |
| Remaining interfering signals (MAG) |  |  |  |  |
| Method | Median fT | Post-hoc comparisons, $p$ | | |
| SCA | 0.000 | SCA | ICA | PCA |
| ICA | 4.686 | $<10^{-19} *$ | | |
| PCA | 16.570 | $<10^{-62} *$ | $<10^{-46} *$ | |
| Original | 35.044 | $<10^{-93} *$ | $<10^{-93} *$ | $<10^{-75} *$ |
| Remaining interfering signals (GRAD) |  |  |  |  |
| Method | Median fT/cm | Post-hoc comparisons, $p$ | | |
| SCA | 0.585 | SCA | ICA | PCA |
| ICA | 1.503 | $<10^{-12} *$ | | |
| PCA | 4.337 | $<10^{-54} *$ | $<10^{-35} *$ | |
| Original | 8.442 | $<10^{-93} *$ | $<10^{-92} *$ | $<10^{-77} *$ |

**Figure 10.**
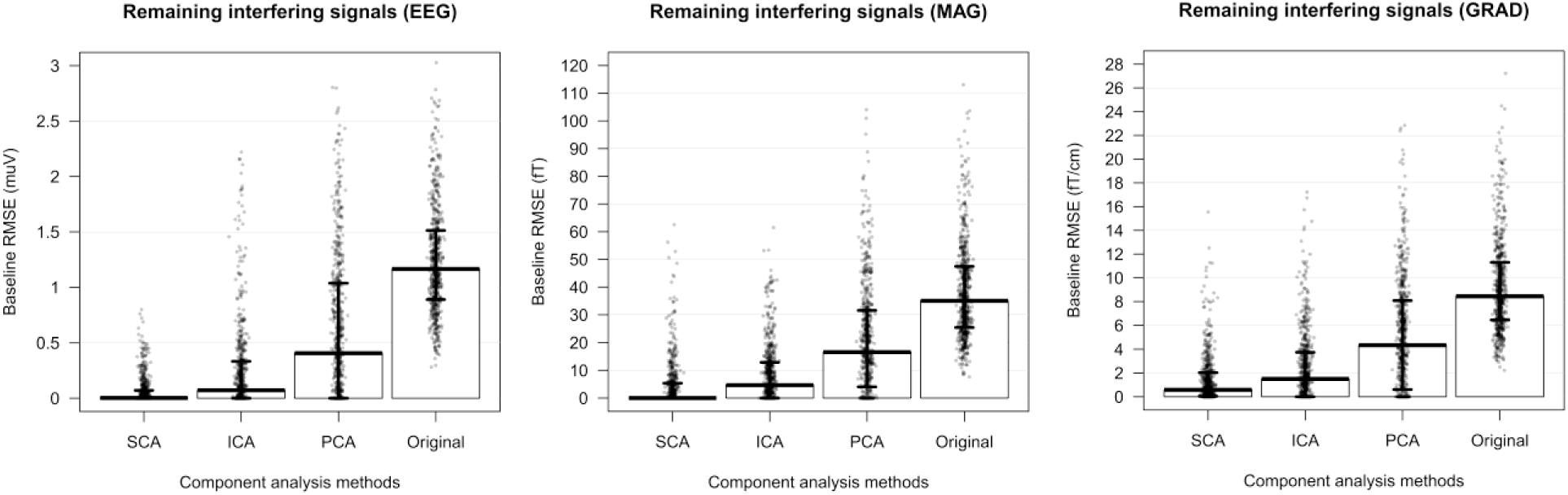
Remaining interfering signals overlapping with component isolated with SCA, ICA, and PCA. Showing the root-mean-squared error (RMSE) from a perfect MEG/EEG waveform baseline of values 0 in the time range surrounding the component of interest, *t*_*comp*_, for EEG, MEG magnetometer (MAG) and MEG gradiometer (GRAD) when applying the SCA, ICA, PCA or the original waveforms.

**Figure 11.**
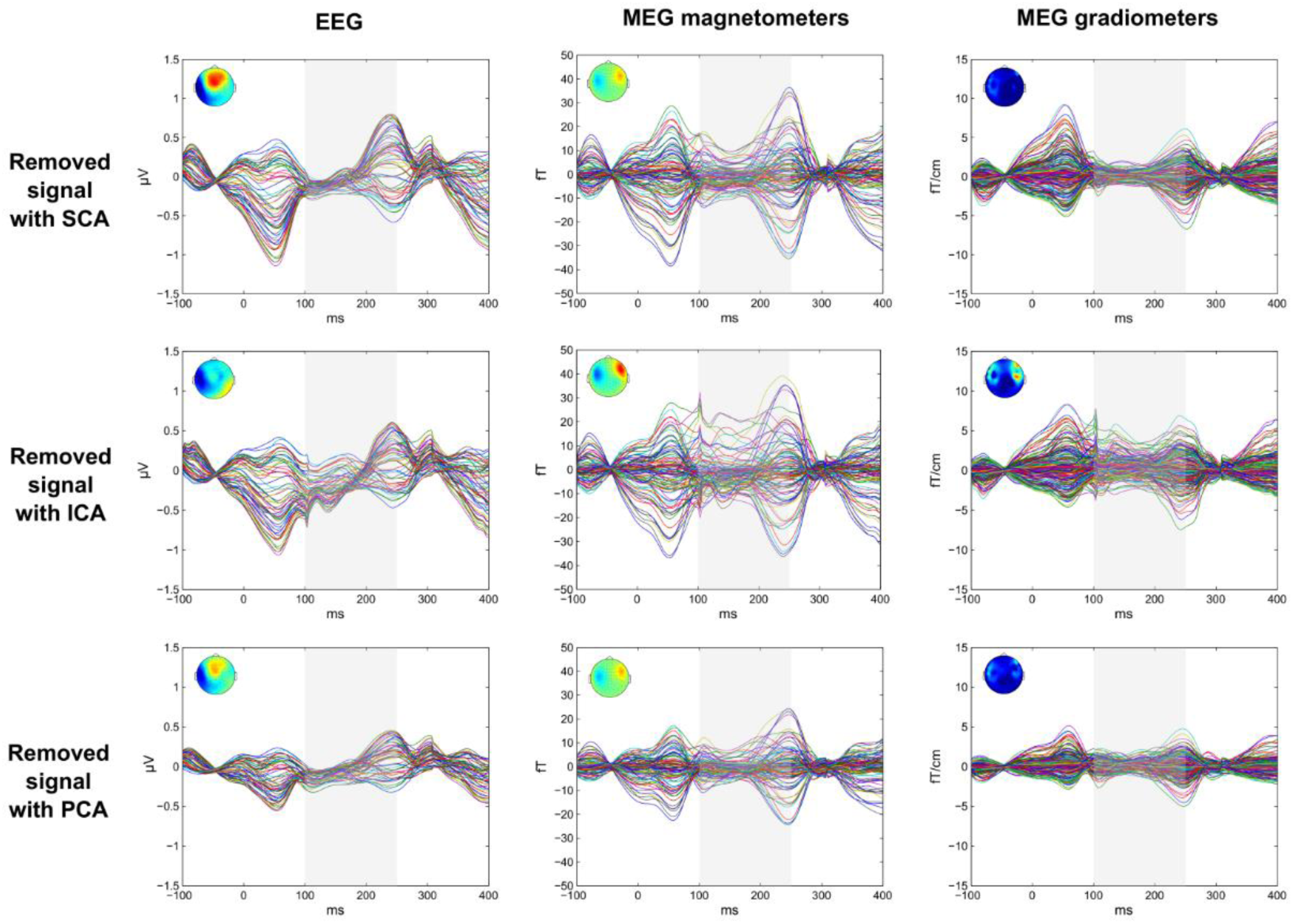
Removed signals with SCA, ICA and PCA. The top row shows the butterfly waveforms for signals removed with SCA for EEG (left), MEG mag. (middle), and MEG grad. (right), the middle row shows the same for ICA and the bottom row for PCA (i.e. the difference waveforms between the original waveforms and extracted component waveforms). Next to each waveform is shown the signal topography in the component of interest time range, *t*_*comp*_.

**Figure 12.**
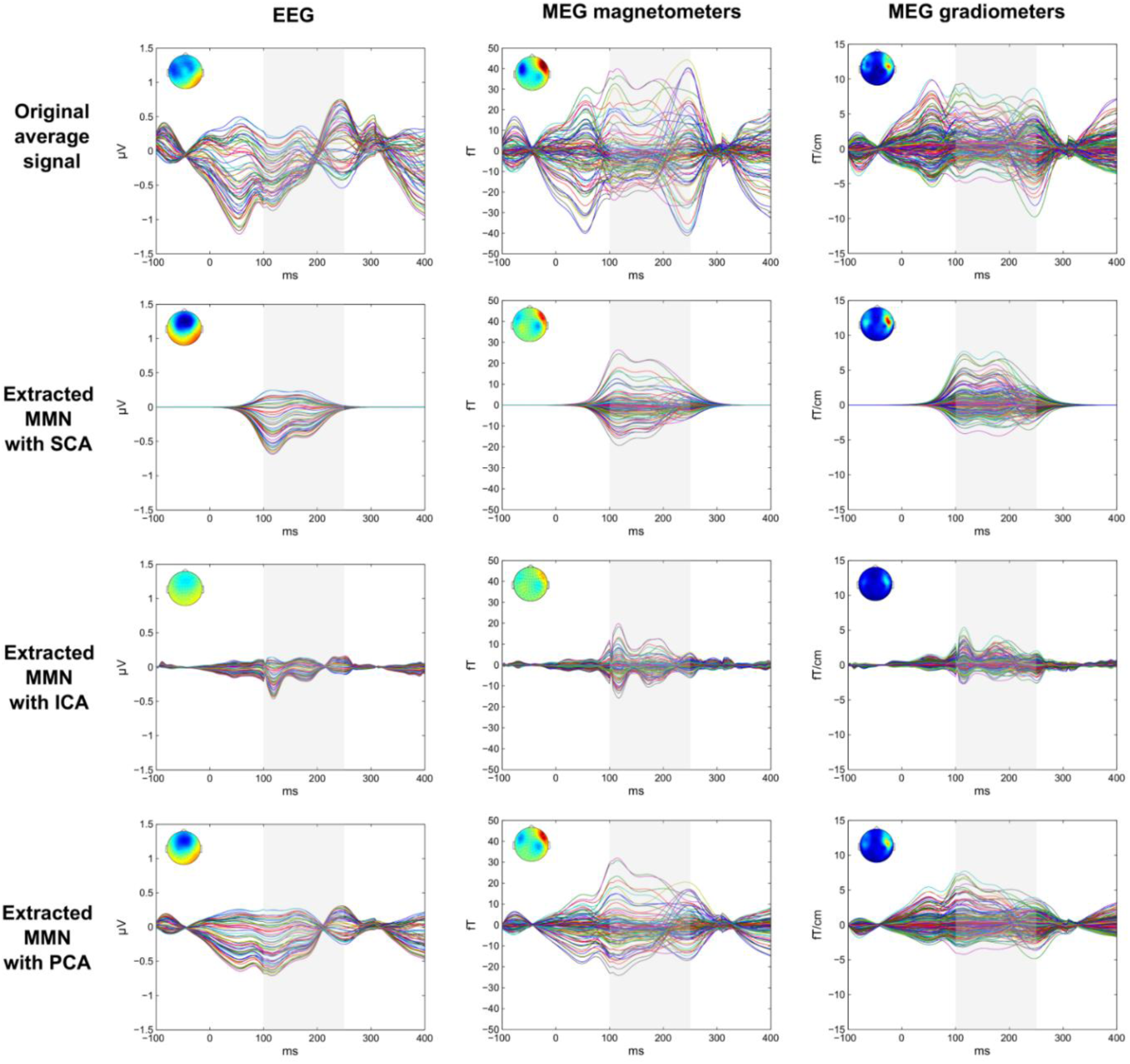
Waveforms extracted with SCA, ICA and PCA. The top row shows the butterfly waveforms for the original average signal for EEG (left), MEG mag. (middle), and MEG grad. (right), the second row shows the same for SCA, the third row for ICA, and the fourth row for PCA. Next to each waveform is shown the signal topography in the component of interest time range, *t*_*comp*_.

## Interim discussion 2

The results of Study 2 show as hypothesized that the novel SCA decomposition method accurately isolates an evoked response of interest, in this case the MMN, from other interfering neural sources in the single-subject MEG and EEG waveforms. In terms of the accuracy in the isolation of the evoked response of interest it is observed that the SCA method clearly outperforms the ICA and PCA methods. Also, the findings show that the evoked response of interest is more accurately represented in the extracted SCA components than in the original measurements. Moreover, as expected, the ICA decompositions suffered in particular from inaccurate representations of the MEG/EEG waveforms. Furthermore, as expected, the PCA components were only partially separated, with interfering signals partially mixed with the component of interest in the PCA decomposition.

Another important finding is that the evoked response of interest is represented by only a few SCA components (1-2 SCA components for the EEG and MEG magnetometers and 1-6 SCA components for the MEG gradiometers), which is crucial for the manual inspection of the components, where the large splitting of the component of interest across a large number of ICA and PCA components causes difficulties with the correct identification of the components of interest with manual inspection. The relatively low number of SCA EEG components containing the MMN component are expected for the relatively high difficulty of in the discrimination of the subtle changes in the stimuli in the applied experimental paradigm, which normally results in low SNIR (Cong et al., 2010), and would thus frequently violate the SCA assumption 1 (SNIR>1) for the EEG. Though, the frequent cases of low numbers of SCA EEG components containing MMN is correctly reflecting the frequent cases with low MMN component amplitudes measured with the EEG. The higher number of SCA MEG gradiometer components containing the MMN might be caused by the higher SNIR and spatial specificity of the MEG gradiometers compare to the EEG and MEG magnetometers (Hämäläinen et al., 1993). Thus, the SCA MEG gradiometer decompositions might more frequently contain additional MMN-related components, which is in line with SCA assumption 2 (components differ in time, width across time or topography, as defined in the methods section).

While Study 1 and Study 2 showed that the SCA method can be applied to isolate a specific component of interest in single-subject measurements, it remains to be verified whether SCA is a reliable method for the study of inter-individual differences measurable in a specific component. In particular, we tested whether the previous findings of increased MMN amplitude to pitch and slide deviants measured in the MEG gradiometers in subjects with higher traits of depression (Bonetti et al., 2017) could be replicated with the SCA method.

## Results of Study 3

In this study we tested SCA on the same dataset as in the previous studies and as in (Bonetti et al., 2017) and categorized the subjects into three groups according to assessed low, medium and high depression trait.

The same effects of individual depression trait on the MMN amplitudes for the spectral features were observed in both the original data and in the MMN components extracted with the SCA method (Table 4). Increasing depression scores were related to increasing MMN amplitudes in response to two spectral changes (pitch and slide) in acoustic features of the sound stimulation (Table 4 and Figure 13 and Figure 14).

**Table 4.**
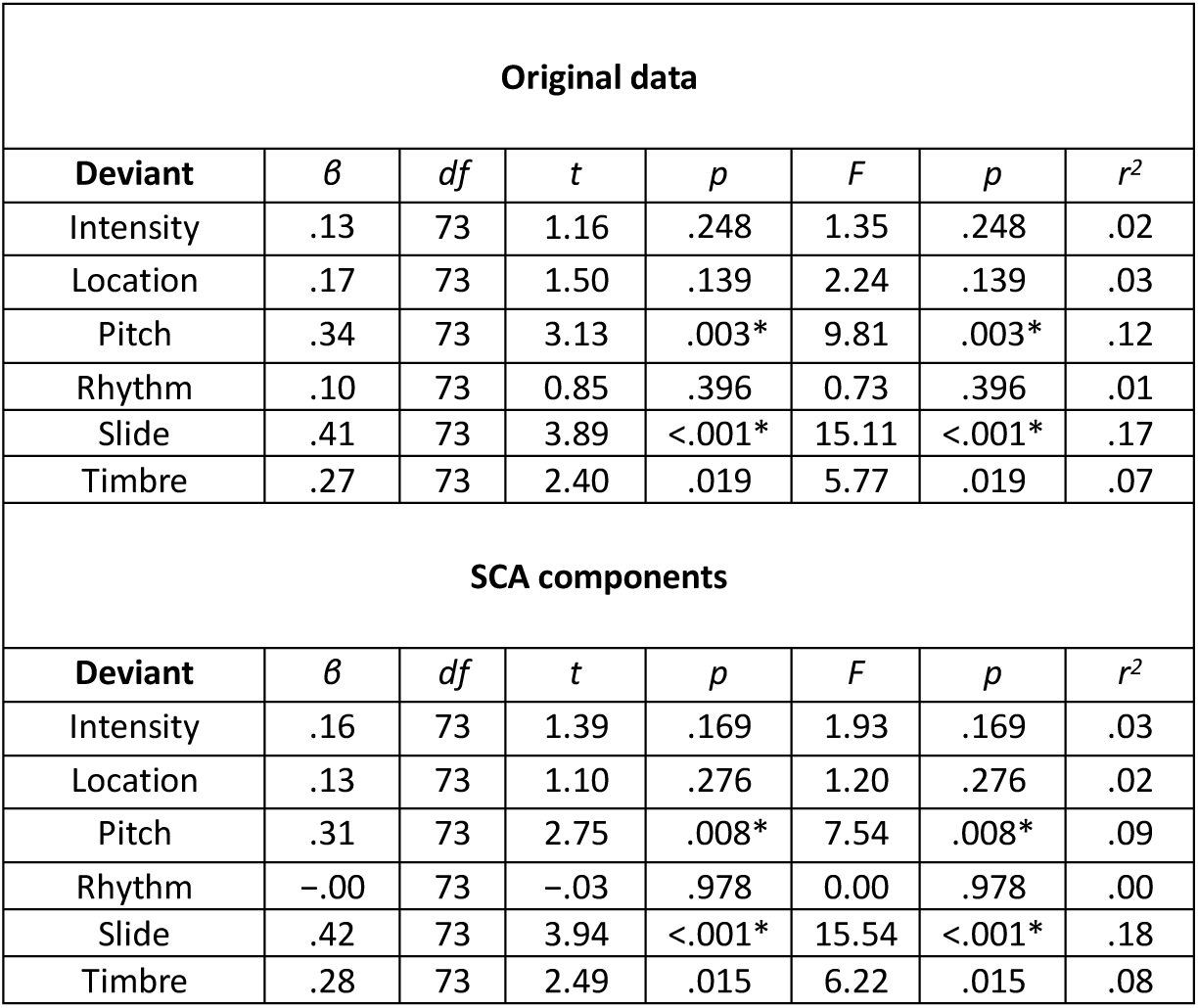
Results of linear regressions between depressive traits score (MADRS) and amplitudes of MMN responses to each of six types of stimulus deviants.

**Figure 13.**
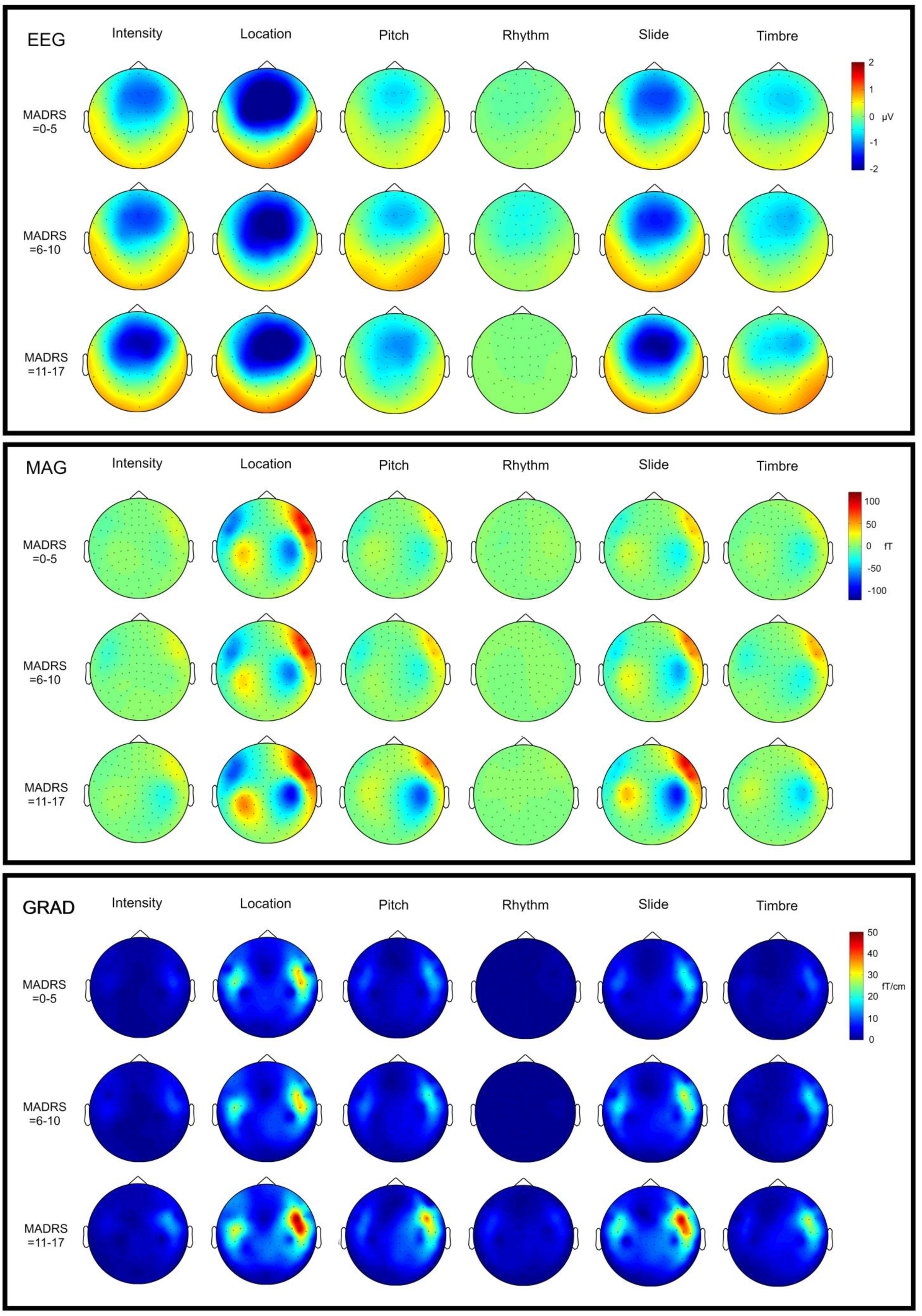
Topographies of extracted MMN response with the SCA method for subjects with low (0-5), medium (6- and high (11-17) depression scores (MADRS) for each type of stimulus deviant.

**Figure 14.**
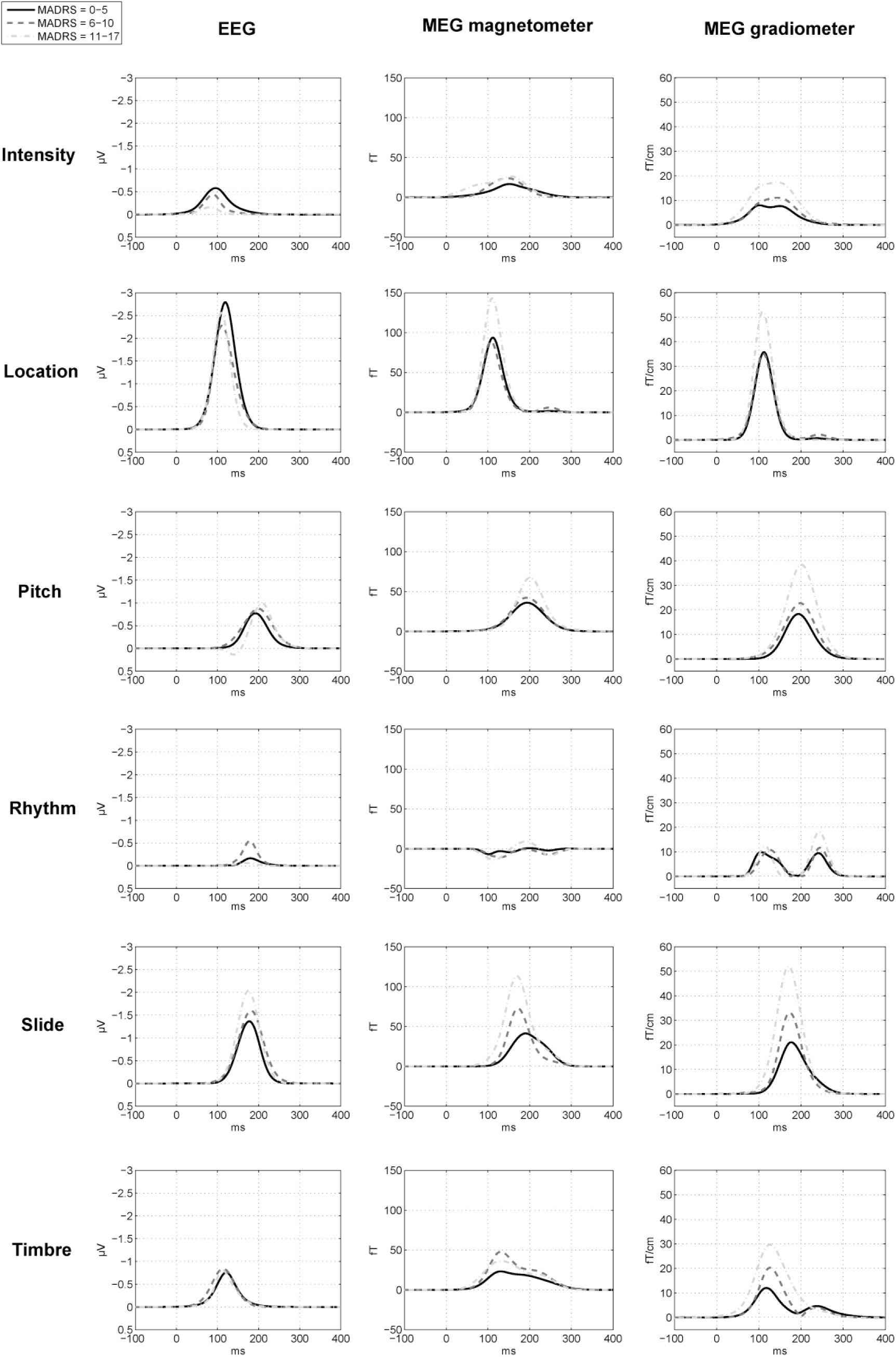
Waveforms of extracted MMN response with the SCA method for subjects with low (0-5), medium (6-10) and high (11-17) depression scores (MADRS) for each type of stimulus deviant.

## Discussion

We here presented a new theory suggesting that the large-scale, evoked or oscillatory, activity in neural networks measured with MEG/EEG can be described by spike timing probability density functions. In the first study we show that Gaussian probability density functions consistently and with high-accuracy model components originating from the brain obtained with MEG and EEG measurements, while the Gaussian functions were unable to model artifact signals, which do not originate from the brain. The results of the second study show that the Gaussian probability density functions can be applied to isolate a specific component of interest, and it is found that the isolated component of interest is represented with higher accuracy with the SCA method than in the original MEG/EEG waveform and in ICA and PCA decompositions. Finally, in the third study we show that the modeling of MEG/EEG components with Gaussian probability density functions is a reliable method for the analysis of inter-individual differences in evoked brain responses relevant to clinical diagnosis. The findings from the three studies presented here suggest that the introduced spike density component analysis (SCA) method offers a novel standard of high-accuracy MEG/EEG analysis.

Our findings support the theory that spiking behavior in certain groups of neurons appears to be systematically stochastically distributed across time, suggesting that this stochastic nature might originate from internal organizations in the brain or from external exposure to environmental stimuli. With regards to brain organization, it has been suggested that the stochastic spike timing might result from the structure of the dendrite pathways in neural networks (Stein et al., 2005; Teramae and Fukai, 2014). Moreover, it has been considered that the stochastic spike timing might originate from variance in environmental stimuli, e.g., the typically Gaussian-shaped timing of the photons reaching the eyes, due to the uncertainty principle in the quantum mechanics of the photon, stimulating the photoreceptors (Pirenne, 1958; Stein et al., 2005), and the Gaussian-shaped Brownian motions within the cochlear of the ears stimulating the auditory nerve fibers (Harris, 1968; Corey and Hudspeth, 1983; Stein et al., 2005). However, the uncertainty in the timing of photons have been shown to be on a scale of nanoseconds (Storzer et al., 2006) and does not seem to affect the timing of photoreceptors (Baylor et al., 1979), and the neural responses to sounds in the brainstem show timing uncertainties ∼1 ms (Don et al., 1977; Bidelman, 2011; Lehmann et al., 2015). These considered external sources of noise therefore seem to be inappropriate explanations for the relatively larger standard deviations in spike timing measured in tenths of milliseconds in the cortex.

Alternatively, the stochastic spike timing behavior in specific brain regions might introduce certain functional advantages over non-stochastic spike timing for the internal processing of certain types of stimulus features. In particular, the function of learning, and the perceptual and cognitive abilities achieved through learning, is assumed to be implemented in the neural networks through spike-timing-dependent plasticity (STDP) (Mcnaughton et al., 1978; Levy and Steward, 1979; Barrionuevo and Brown, 1983; Stein et al., 2005; Lord et al., 2017). This means that for a general Hebbian type of learning to occur, it is necessary that the spikes, which reflect the neurotransmission process, between two or more neurons in a neural network are overlapping in time. For Hebbian learning based on neurotransmission from lower level areas to be integrated across time, e.g. auditory patterns or visual movements, the STDP in higher level association areas would improve with a stochastic spike timing function for systematically increasing the overlap of the spikes. This explanation is consistent with the observations of larger spike timing distributions widths in cortical association areas (Picton et al., 1974) compared to more narrow spike timing distributions widths observed in the brain stem (Picton et al., 1974), in primary somatosensory cortex (Forss et al., 1994), and in specialized motor and cognitive networks comprising regions of the frontal lobe, basal ganglia and cerebellum (Kelly and Strick, 2003; Stein et al., 2005), where the last regions are functionally specialized in fast processing and accurate timing (Dreher and Grafman, 2002; Bostan et al., 2010). In future studies it remains to be investigated whether, for example, stimuli that require integration of features across longer time windows evokes cortical responses with larger standard deviations across time compared to stimuli that requires fast processing and high timing accuracy. Moreover, it would be interesting to see whether the widths of the spike timing distributions in motor neurons vary dependent on the type of action.

Until now we have mainly considered MEG/EEG analyses in the time domain. Based on the present theory and findings of low explanatory power of narrowband sinusoids for evoked MEG/EEG signals, it could be considered that the observations of cross-frequency couplings in the frequency domain (Lakatos et al., 2005; Buzsaki et al., 2012) might reflect that high-frequency sinusoids are phase-locked to the slower sinusoids, because the lower and higher frequency sinusoids conjointly describe the shape of an underlying non-sinusoidal broadband component. While most MEG/EEG analyses in the frequency domain focus on narrowband oscillations (Pfurtscheller and da Silva, 1999), in modern signaling theory investigations of signals from sine waves to square pulses have found that the Gaussian function provides an optimal signal shape, which allows a certain amount of timing uncertainty in a communication system (Turletti, 1996). According to the Gaussian minimum-shift keying (GMSK) scheme in signaling theory, the Gaussian function provides an optimal compromise between minimization of the overlap in time and of the occupied frequency bandwidth (related by σ(frequency domain) = 1/σ(time domain)) (Turletti, 1996). Also, a discussion has recently been introduced specifically concerning Gaussian shapes in MEG/EEG power spectra (Haller et al., 2018). The here presented theory of large-scale stochastic neuronal spike trains and the SCA method could provide a theoretical framework and method for estimating the power, peak frequency, and bandwidth and topography of the Gaussian shaped SCA components in the frequency domain.

Furthermore, in source location analysis, PCA and ICA has been applied for separating the overlapping components to improve the source location modeling (Vigario et al., 2000; Zhukov et al., 2000; Richards, 2004; Tsai et al., 2006; Reynolds and Richards, 2009). The findings obtained here showed that the SCA method more accurately isolates a component of interest than PCA and ICA, and it is likely that more accurate source reconstructions can be achieved by combining SCA with source location analysis. The combination of SCA and source location analysis might also offer interesting possibilities for simplifying the multichannel MEG/EEG data. For example, the description of a component could be reduced to nine parameters: three Gaussian parameters, three location parameters, and three orientation parameters, which in comparison to a complete multichannel waveform in a range of around hundred channels multiplied by more hundreds time samples offer better possibilities for MEG/EEG data sharing and meta-analysis by simplifying the data.

## Conclusion

We introduce a theory suggesting that the large-scale stochastic spiking activity observed in MEG/EEG measurements can be accurately described by probability density functions. Findings from our three studies mutually supported the theory, and the findings suggest that a novel standard of high-accuracy MEG/EEG analysis can be achieved with an introduced spike density component analysis (SCA) method. The theory and findings presented here offer a novel standard of high-accuracy MEG/EEG analysis, which is of particular relevance to the investigations on individual differences in brain function and single-subject clinical diagnoses.

## Methods for Study 1

### Repository dataset

A pre-existing dataset was used consisting of 564 average ERP/ERF waveforms recorded from 94 human subjects each exposed to six different experimental conditions under the “musical multi-feature no-standard” stimulus paradigm and recorded with 366 channel simultaneous EEG (60 electrodes) and MEG (102 axial MEG magnetometers, and 204 MEG planar gradiometers) at the Biomag Laboratory of the Helsinki University Central Hospital (for further details, see e.g. (Bonetti et al., 2017)). The dataset was a part of the data repository obtained under the research protocol named “Tunteet”, approved by the Coordinating Ethics Committee of the Hospital District of Helsinki and Uusimaa (approval number: 315/13/03/00/11, obtained on March the 11th, 2012). Findings based on this dataset have previously been published in studies on noise sensitivity (Kliuchko et al., 2016), comparison of artifact corrections methods (Haumann et al., 2016), relationship between MMN amplitude and depressive traits (Bonetti et al., 2017) and working memory skills (Bonetti et al., 2018).

### Data preprocessing

MEG data was preprocessed with Elekta Neuromag™ MaxFilter 2.2 Temporal Signal Space Separation (tSSS) (Taulu and Hari, 2009) (with automatic detection and correction of bad MEG channels; default inside expansion order of 8; outside expansion order of 3; automatic optimization of both inside and outside bases; subspace correlation limit of 0.980; raw data buffer length of 10 seconds). Afterwards MEG and EEG data was further preprocessed with the FieldTrip version r9093 toolbox for Matlab (Donders Institute for Brain, Cognition and Behaviour/Max Planck Institute, Nijmegen, the Netherlands) (Oostenveld et al., 2011) and Matlab R2013b (MathWorks, Natick, Massachusetts). EEG and MEG waveforms were bandpass filtered between 1-30 Hz. EEG channels were inspected and bad channels corrected them with interpolation of the neighboring channels. Eye blink and EKG artifacts were inspected and corrected with ICA (Makeig et al., 1996).

### Extraction of average ERP/ERF waveforms

Trials were extracted and ERP/ERF waveforms averaged for six experimental conditions consisting of (1) intensity, (2) location, (3) pitch, (4) rhythm, (5) slide and (6) timbre deviants, and average ERP/ERF waveforms for a standard condition was subtracted to obtain the evoked mismatch negativity (MMN) waveforms (for further details, see (Bonetti et al., 2017)). A duplicate of the same average evoked MMN waveforms with presence of external artifactual signals was created by excluding the preprocessing procedures, before the trials and average MMN waveforms were extracted from the same dataset.

### Spike density component analysis

Assuming that MEG/EEG waveforms in the time domain can be modeled with Gaussian functions (Formula 1), the *α* parameter describes the maximum spike rate, which is equivalent to the amplitude of a component in the MEG/EEG waveform (Table 5). The *μ* parameter denotes the expected latency, corresponding to the latency of a component in terms of conventional MEG/EEG waveform analysis, while the *σ* parameter defines the spike timing uncertainty, or width of a component in the MEG/EEG waveform (Table 5).

**Table 5.**
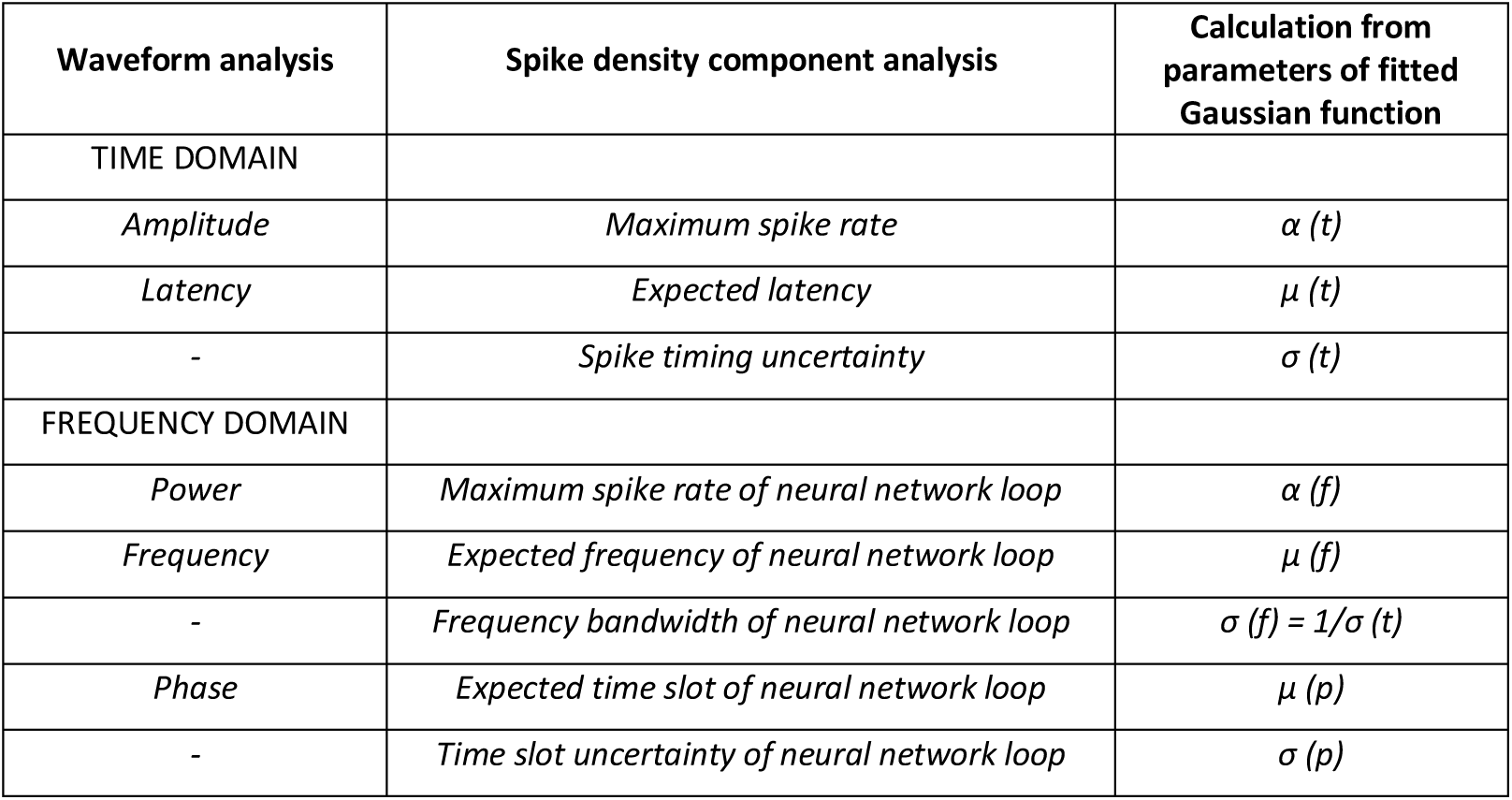
Relationships between spike densities and MEG/EEG waveforms.

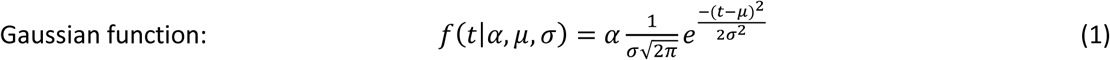

The Gaussian function applied in the time domain analysis will result in another Gaussian function applied in the frequency domain analysis (Bracewell and Bracewell, 1986). In the frequency domain, the *α* parameter describes the maximum spike rate of a neural network loop, equivalent to the observed power of the oscillation (Table 5). The *μ* parameter in the frequency domain defines the expected frequency of the neural network loop, and the *μ* parameter in the phase domain defines the expected time slot of the neural network loop (Table 5). The uncertainties in frequency and phase are represented by the *σ* parameters for frequency and phase (Table 5).

Another function often considered in spike density analysis is the gamma function (Formula 2) (Gerstein and Mandelbrot, 1964; Barbieri et al., 2001; Maimon and Assad, 2009). Here the shape parameter, *k*, defines the regularity of the spike timing, where higher value of *k* denotes more regular spike timing, which approaches a Gaussian distribution, while lower value of *k* denotes more random and skewed spike timing, differing from a Gaussian distribution (Maimon and Assad, 2009). For example, it has been found that neurons in the rhesus monkey higher level visual association area show more regular spike timing, *k*≈8, compared to the more random and skewed spike timing of neurons in the lower level visual areas, *k*<5 (Maimon and Assad, 2009).

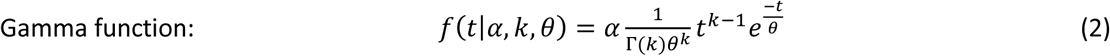

Moreover, we suggest investigating the sine function, which is the foundation for analyzing narrowband oscillations in MEG/EEG waveforms, the theta, alpa, mu, beta and gamma waves (Formula 3) (Pfurtscheller and da Silva, 1999). It should here be noted that the sine function reflects regular changes in the spike rate related to neural network loops involving polarity reversals (not to be confused with the frequency of the spike rate).

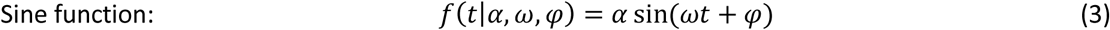

Spike density component analysis (SCA) in the time domain is performed on each average ERP/ERF waveform by following an automatized iterative procedure (Figure 15) (example outputs are created with an open source FieldTrip-compatible Matlab function freely available at [Github upon publication] for decomposing any MEG/EEG/ECoG/iEEG data into SCA components). With SCA it is assumed that:

**Figure 15.**
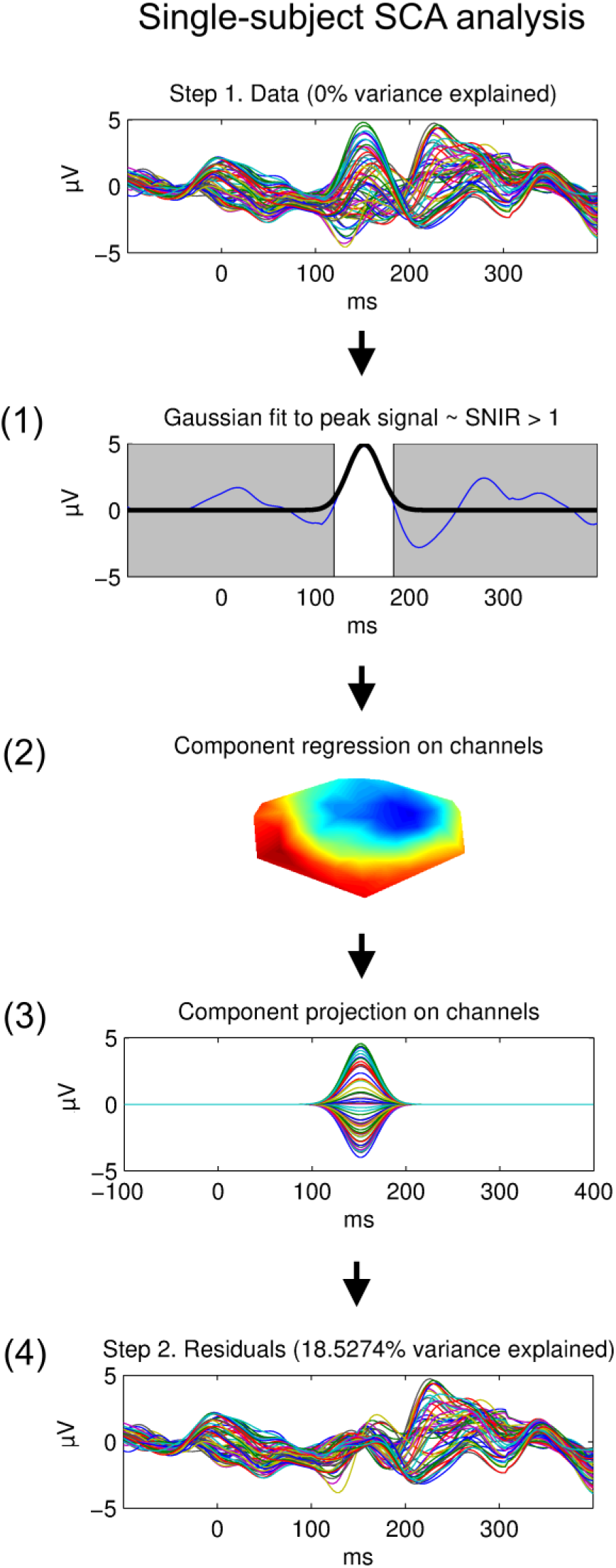
Single-subject SCA analysis. Showing the iterative four-step sub-routines of the SCA algorithm.

1. Components exists at signal-to-noise and interference ratios (SNIR) > 1. (SNIR here refers to background noise and artifacts, not overlapping components from the brain).
2. Components differ in time, width across time or topography.

The SCA decomposition procedure is similar to PCA, though, since SCA finds components with specific temporal shapes, each SCA step begins by finding the maximum amplitude across channels and time (instead of finding the maximum variance across the multichannel waveforms). First, the Gaussian function parameters (Formula 1 and Figure 1 bottom) are estimated with the *fit.m* Matlab function, in the part of the waveform of the channel with maximum amplitude, on the time samples that are estimated to be valid with respect to the SNIR > 1 assumption, which are found by extending the time samples around the peak amplitude time sample until the nearest valley or baseline crossing is reached (Figure 15 (1)). The SCA component waveform is modeled by applying the fitted function parameters. Since part of the data might contain non-Gaussian signals, if the Gaussian function parameter estimation fails, i.e., the errors between the modeled and measured data exceed the 95% confidence intervals, the raw curve within the time samples is applied as a substitute instead of a modeled waveform, while the search for Gaussian shapes of lower amplitudes continues in the subsequent iterations.

Second, the component weighting matrix, *W*_*n,c*_, for the weighting of each component waveform, *n*, on each channel, *c*, i.e. the topography each component (Figure 15 bottom), is estimated with linear regression of each component waveform, *x*_*n*_, on each residual channel waveform, *y*_*c*_, based on the formula *y*_*c*_ = *W*_*n*,*c*_*x*_*n*_, with the Matlab function *mldivide.m* (Figure 15 (2)). To minimize the influence of false partial correlations between the component and channel waveforms, the linear regression is based on the complete range of time samples.

Third, the modeled component waveform is multiplied by the channel weight vector, *W*_*n,c*_, to create a projection of the component waveform on the channels (Figure 15 (3)). Fourth, the component waveform projected back on the channels is subtracted from the multichannel waveforms to obtain the residual waveforms (Figure 15 (4)).

The SCA procedure is performed, and components are estimated iteratively, as long as the subtraction of the estimated component waveform projected on the channels results in a decrease in the sum of the residual waveforms across channels and time. When the sum of the residual waveforms across channels and time increases or reaches a value of 0, the SCA algorithm stops.

SCA based on gamma functions was performed using the exact same procedure, except that the parameters of the gamma function (Formula 2) were estimated with the *gamfit.m* function in Matlab. SCA with sine functions was also achieved with the exact same procedure, although the parameters of the sine function (Formula 3) was fitted only to the sine arc, or sine half-wave, with the *fit.m* Matlab function, and only the sine half-wave was applied in the back projection of the components onto the channel waveforms.

### Performance evaluation

The performance of the algorithms was evaluated by measuring the explained variance of the measured ERP/ERF waveform by the modeled SCA waveforms, based on the mean square of the Pearson‘s product-moment correlation coefficients between each modeled and measured waveform across the channels. In addition, the peak amplitudes (across the complete time range) were obtained from the residual waveforms, showing the peak amplitudes in the part of the waveform that could not be modeled by the SCA components (including any component substitutes with raw curves applied during the SCA procedure).

### Statistical analysis

The performance evaluations showed general tendencies towards high performances, resulting in skewed performance distributions diverging from normality in the positive direction (Kolmogorov-Smirnov and Shapiro-Wilk tests shows overall violations of the normality assumption at *p*<.001). Therefore, differences in performance, as defined in the preceding section, were tested with Friedman’s ANOVA by ranks, and post-hoc comparisons were conducted with Wilcoxon signed rank tests.

## Methods for Study 2

### Repository dataset

The repository dataset for Study 2 was the exact same as in Study 1.

### SCA, ICA and PCA decomposition

The SCA decompositions were performed following the same procedure, except that a few minor improvements for increasing the robustness of the SCA modeling were included:

1. The bias from interfering signals in the baseline was minimized with a baseline correction to the median (instead of the mean) values in the time frame of −100 to 0 from stimulus onset.
2. The bias from interfering signals in the MEG/EEG waveform (in the time samples distant from the peak amplitude approaching the valleys or baseline, where the amplitude increases for the background noise and overlapping components in relation to the component of interest) on the Gaussian function parameter estimates was reduced. This was achieved by estimating the Gaussian function parameters with the bisquare weighting function for iteratively reweighted least square error (LSE) (with the standard tuning parameter value 4.685) (Holland and Welsch, 1977) in the Matlab *fit.m* function (instead of conventional LSE estimates).
3. The bias in the spatial domain from overlap of interfering signals across MEG/EEG channels was minimized. This bias minimization was implemented by applying linear regressions for estimating the component projection weights (*W*) performed with the bisquare weighting function for iteratively reweighted least square error (LSE) (with the standard tuning parameter value 4.685) (instead of conventional LSE).

The SCA results were compared with those of principal component analysis (PCA) and independent component analysis (ICA).

The PCA decomposition was performed by applying the *pca.m* Matlab function. PCA is an iterative procedure by which the waveform explaining most of the variance in the data is estimated and subtracted from the data, while subsequent components explaining most of the remaining variance in the data are repeatedly estimated (Jung et al., 1998). A constraint is imposed that each weaker component must be topographically orthogonal to the preceding component, in order to increase spatial independence between the components (Jung et al., 1998). While PCA typically succeeds suppressing the signal of spatially dissimilar components explaining less variance from components explaining more variance, PCA fails in separating mixed signals from spatially similar components (Jung et al., 1998).

ICA achieves higher spatial accuracy than PCA by separating the mixed multichannel signals into spatially statistically independent sources across time (Groppe et al., 2008). However, weaknesses of ICA concern its dependency on estimating the projection weights based on statistics obtained across time, whereby its accuracy decreases with lower SNR (related to higher background noise) (Comon, 1994) and fewer time samples (Jung et al., 2000). Therefore, ICA is able to isolate, e.g., eye blink artifact components accurately, because they exhibit high SNR and can be estimated based on several time samples in the continuous MEG/EEG recording (Haumann et al., 2016). However, the assumptions underlying ICA are violated for brain responses that show either low SNR in the continuous recording, or few time samples if signal averaging is applied to increase the SNR. Therefore, it is likely that the evoked response waveforms will be distorted when ICA is applied to decompose these signals (Groppe et al., 2008). Also, if ICA estimates are attempted to be estimated from concatenated averages of MEG/EEG signal across experimental conditions and subjects, this results in summary statistics, which ignores the individual variance across conditions, which compromises the single-subject analysis (Groppe et al., 2008).

The ICA decomposition was performed with the Infomax algorithm, implemented in the *runica.m* function for Matlab, which has been shown to be among the most accurate ICA algorithms for EEG data (Delorme et al., 2007; Crespo-Garcia et al., 2008) and is also commonly applied for artifact correction for MEG and EEG data (Haumann et al., 2016). First, the rank of the average ERP/ERF waveform was estimated with the *rank.m* Matlab function. The resulting rank number was given as input to the *runica.m* function for the initial PCA-based dimensionality reduction step prior to the ICA procedure. In cases where the ICA decomposition resulted in imaginary numbers in the component waveforms or topographies, due to overestimates of the rank, the assumed rank and PCA-based dimensionality output was reduced by 1, and the ICA decomposition repeated, until the resulting ICA estimates contained only real numbers.

### Automatic component of interest extraction based on template match

The stimulus paradigm was specifically designed to evoke mismatch negativity (MMN) responses, and the investigated dataset contained a total of 1692 cases of averaged MEG/EEG multichannel waveforms with MMN responses to be analyzed (564 cases simultaneously recorded with EEG, MEG magnetometers and MEG gradiometers). Since each case was analyzed with SCA, ICA, and PCA, the resulting set of 5076 decompositions in total was relatively large for conventional manual identification and extraction of the MMN components. Moreover, it was important to ensure that the MMN components were extracted following the exact same procedure across the SCA, ICA and PCA decompositions. Therefore, an automatic component of interest extraction procedure was developed, which was based on a template matching approach (for similar methods, see (Lee et al., 2003). Since the dataset contained six different types of deviant stimuli affecting the MMN component, all MMN components were identified separately for each type of deviant stimulus. The grand average group-level MEG/EEG waveforms were applied as templates and matched against the SCA, ICA and PCA components.

First, the match between each component and the group-level template was calculated in terms of Pearson‘s correlation *r*-estimates. Reliable time points, *t*_*comp*_, containing the MMN component waveform in the template were isolated by finding the peak amplitude, and in the waveform of the channel with the peak amplitude extending the selected time points around the peak amplitude until they reach the baseline. In addition, *t*_*comp*_ was constrained to be within the typical MMN component range of 75-250 ms. A template topography vector was calculated as the mean channel values across the time window *t*_*comp*_. Also, component topographies were calculated for each component as the mean projected channel values across the time window *t*_*comp*_. Next, topography *r-*values, *r*_topo_ were estimated for each component by correlating each component topography with the template topography. Moreover, waveform mean *r*-values, *r*_wave_, were estimated for each component, in the time window *t*_*comp*_, by correlating the component waveform projection on each channel with the template waveform for the channel and deriving the mean *r*-value across channels. Based on this, a pseudo *R*^*2*^-value, taking into consideration the importance of the match in component polarity, topography and waveform, was estimated for each component as *R*^2^ = *polarity* × |*r*_*topo*_| × |*r*_*wave*_|, where the *polarity* is −1 if either *r*_topo_ or *r*_wave_ is negative and else +1. Any components with *R*^*2*^<0 were excluded from further analysis. For each SCA, ICA, and PCA decomposition, the single-subject components were sorted in descending order according to their *R*^*2*^-values indicating their match with the group-level template component.

Finally, following the order of the *R*^*2*^-values, each component was projected and summed into the extracted channel waveforms, as long as the addition of a component resulted in an increase in the correlation, *r*, which was initially set to *r*=0, and which was calculated by correlating the projected component waveform with the template waveform in the time window *t*_*comp*_ and obtaining the mean *r* across channels.

### Performance calculations

The accuracy of the SCA, ICA, PCA methods for decomposing MMN components and the accuracy of the original MEG/EEG waveforms in representing the MMN components was evaluated and compared. First, the number of sub-components representing the MMN with SCA, ICA and PCA was counted. The accuracy of the MMN topography was calculated as the *r*^2^-value based on the squared correlation coefficient between the extracted MMN component topography and the group-level MMN topography within the time points *t*_*comp*_. Also, the accuracy of the MMN waveform was calculated as the mean *r*^2^-value equal to the mean squared correlations coefficients between the extracted single-subject MMN waveform and the group-level MMN waveform within the time points *t*_*comp*_, and the mean *r*^*2*^ was derived across channels. In addition, the ability to remove interfering signals was evaluated by calculating the root-mean-squared error between the ideal baseline with values of 0 and the waveform values outside the MMN time range *t*_*comp*_.

### Statistical analysis

Since the performance values were not normally distributed (most Kolmogorov-Smirnov and Shapiro-Wilk test results are *p*<.001), differences in performance were, as in Study 1, tested with Friedman‘s ANOVA by ranks, and post-hoc comparisons were conducted with Wilcoxon signed rank tests.

## Methods for Study 3

### Repository dataset

For Study 3 the same dataset was applied as in Study 1 and Study 2 and in the a previously reported study on effects of depressive traits on MMN (Bonetti et al., 2017). The study included a subset of 75 subjects rated on the Montgomery–Åsberg Depression Rating Scale (MADRS) (Bonetti et al., 2017).

The measured MMN components were categorized according to six types of auditory deviants that evoked the MMN: 1) intensity deviants with −6 dB change in sound amplitude, 2) location deviants where the sound amplitude in one of the stereo sound channels was lowered, 3) pitch deviants with 1.4% change in tone frequency, 4) rhythm deviants with shortening of a tone by 60 ms, 5) slide deviants with gradual change in tone frequency, and 6) timbre deviants with an “old time radio” sound spectrum filter-effect (Bonetti et al., 2017).

### Statistical analysis

The mean amplitude was measured in a 30-ms time window centered on the peak latency in the grand average, measured separately for each deviant type. As in the previous study the effect was investigated for the MEG gradiometers (Bonetti et al., 2017), and the combined gradiometer channels MEG 1322+1323 above the right hemisphere which showed the largest amplitude was applied for testing with linear regression. Statistical test results were obtained with linear regressions between the MARDS score and the extracted mean MMN amplitude for each type of deviant.

## Acknowledgments

Center for Music in the Brain is funded by the Danish National Research Foundation (DNRF117).

